# Distinct Progression Paths Underlie the Diversity of Pancreatic Papillary Cystic Neoplasms

**DOI:** 10.64898/2026.10.06.757128

**Authors:** Camille Pignolet, L Aguilera Munoz, L Bourlet, T Bourega, L Gomes, D Mendes, N Theou Anton, N Colnot, M Albuquerque, L de Mestier, S Dokmak, A Couvelard, V Rebours, R Nicolle, J Cros

## Abstract

Surgical decisions for intraductal papillary mucinous neoplasms (IPMNs), precursors of pancreatic cancer, depend on distinguishing low-from high-grade dysplasia, a distinction that molecular markers could refine. We microdissected 433 tumor regions from 135 patients and integrated transcriptomics, deep-learning histology, and *KRAS/GNAS* RNA-called variants. RNA programs and morphological motifs separated lesions by phenotype, intestinal versus others, rather than by dysplasia grade. Along gastric IPMN trajectories, low-grade lesions recapitulated the pit-chief polarity of normal gastric glands, whereas high-grade lesions were indistinguishable from pancreatobiliary lesions in morphology, RNA programs, and driver mutations. Intestinal IPMNs arose through two routes: a canonical *GNAS*-associated pathway and a *KRAS*-associated shift from low-grade gastric features, with lesions sharing driver mutations retaining common programs across phenotypes. Resolving these phenotypic trajectories clarified distinct progression paths and enabled phenotype-specific prediction of high-grade dysplasia in mucus aspirated from 94 resected lesions. These findings support a promising fluid-based assessment of IPMN status.

**Statement of significance:** IPMN surgery depends on distinguishing low-from high-grade dysplasia. Combining multiregion microdissected transcriptomics with deep-learning morphology across 135 patients, we resolve intestinal and gastric/pancreatobiliary trajectories supported by shared driver mutations, including a gastric-to-intestinal switch, delineating phenotype-specific progressions. These trajectories can be assessed from pancreatic fluid, which could inform surgical decisions.

## Introduction

Pancreatic ductal adenocarcinoma (PDAC) is one of the deadliest human cancers, with a five-year relative survival of about 13% (1). This dismal prognosis reflects late detection, when curative treatment is no longer possible. Pancreatic cystic lesions are common, with a prevalence of 16% on magnetic resonance imaging (MRI) in the general population (95% CI: 13-18%), rising to 38% after the age of 80 (2). Intraductal papillary mucinous neoplasms (IPMNs) are the most frequent of these lesions. As radiologically detectable precursors of PDAC, they represent a unique opportunity for cancer interception, and their incidental diagnosis is steadily increasing (3). However, the vast majority never progress to malignancy, questioning the role and timing of surgery. Resection carries substantial morbidity, and preoperative risk stratification remains imprecise: only about one third of resected IPMNs harbor high-grade dysplasia, the intended target of resection, the others are resected either too early (30% low-grade dysplasia) or too late (35% invasive carcinoma) (4). Current management relies solely on clinical and radiological criteria, ignoring the underlying biology of these lesions (5,6). Reliable molecular biomarkers predicting malignant progression are therefore critically needed.

IPMNs are histologically classified into gastric, intestinal, and pancreatobiliary (PB) subtypes based on epithelial morphology and mucin expression (7). Gastric-type lesions express *MUC5AC* and *MUC6*, arise predominantly in branch ducts, and carry *KRAS* and *GNAS* mutations in about half of cases (7–9). Intestinal-type lesions express *MUC2* and *CDX2*, more frequently involve the main duct, and are enriched for *GNAS* mutations, while *KRAS* alterations are less frequent than in gastric-type IPMN but still affect a sizable subset (7, 8, 10). They are also clearly distinct from gastric-type lesions at a broader level, with greater genomic instability and divergent transcriptomic and epigenetic profiles (8). PB-type lesions are rarer and less well known. They express *MUC1* and combine the highest malignant risk with frequent *KRAS* and rare *GNAS* mutations (8). A fourth pattern “oncocytic” has since been reclassified as a distinct entity (Intraductal oncocytic papillary neoplasm (IOPN)) in the 2019 WHO classification, defined by *PRKACA/PRKACB* fusions and a near-absence of *KRAS* and *GNAS* mutations (6). Whether IOPN is biologically distinct remains debated, since like *GNAS*-driven IPMN, they converge on cAMP pathway activation. Clinically, however, they are equivalent: both present as cysts, cannot be separated on imaging, and can progress to PDAC. We, like others, therefore refer to them collectively as pancreatic papillary cystic neoplasms (PPCNs) (11). More broadly, histological grading and subtyping can be challenging, as individual lesions may display combinations of distinct epithelial morphologies and architectural patterns (12). Thus, PPCNs can exhibit multiple phenotypic features within the same lesion, raising questions about the biological relationships between these phenotypes and the mechanisms underlying their coexistence. These observations suggest that phenotypic boundaries may be less discrete than implied by conventional histological classification.

IPMN progression is currently most often modeled as a single linear sequence, in which low-grade dysplasia gives rise to high-grade dysplasia and then to invasive carcinoma. This model is reflected in the binary histological grading system, which remains partly subjective and may inform surgical decision-making when available. It also lacks molecular support: no consistent transcriptomic or genomic signature separates the two grades (7,8,13), and although *TP53*, *CDKN2A* and *SMAD4* alterations are enriched in high-grade lesions, a substantial fraction of high-grade lesions carry none of them (14). Grade may therefore be a shared label applied to molecular evolutions that are biologically distinct, depending on the phenotype. We therefore hypothesized that IPMNs follow several phenotype-specific routes of progression rather than a unique one, each with its own high-grade endpoint, and that trans-phenotype transitions are possible, as suggested by biphenotypic lesions. Resolving these routes is a prerequisite for predicting which lesions might progress.

To address these questions, 418 regions from 127 patients were analyzed by a combined and spatialized histomolecular transcriptomic approach (Fig. 1A). These data were used to characterize the morphomolecular landscape of PPCN and investigate progression trajectories, with key observations supported by spatial transcriptomics. Transcriptomic programs identified in PPCN were then compared with established PDAC transcriptomic subtypes to explore potential malignant endpoints. Finally, IPMN mucus-derived RNA from an independent cohort of 94 patients with PPCN was analyzed to determine whether these signatures could predict phenotype-specific dysplasia grade from these minimally invasive samples.

**Figure 1.**
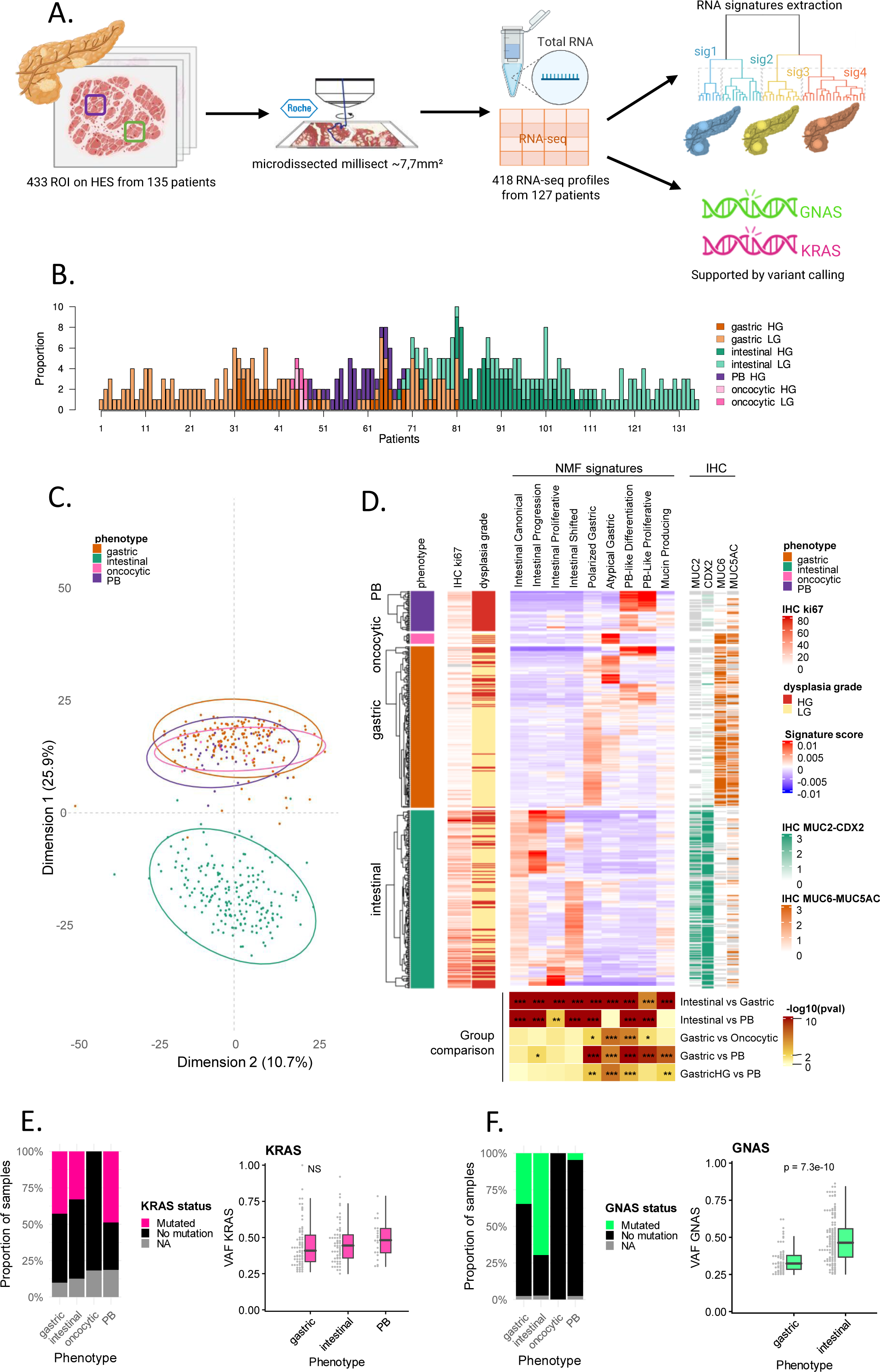
Integrative transcriptomic profiling of IPMN reveals two primary molecular axes. **(A)** Study workflow. Morphologically uniform regions of interest (ROIs) were annotated on hematoxylin-eosin-saffron (HES)-stained sections of FFPE surgical specimens, microdissected, and profiled by bulk RNA sequencing. Expression data were decomposed into transcriptional programs by consensus NMF, and *KRAS/GNAS* hotspot mutations were called from RNA sequencing data. **(B)** Number of ROIs per patient, colored by phenotype and dysplasia grade. Patients contributed one or several ROIs depending on intra-lesional heterogeneity. PB, pancreatobiliary; LG, low-grade; HG, high-grade. **(C)** PCA of RNA sequencing profiles across 418 ROIs, performed on the expression matrix of the 1,000 most variable genes, colored by histological subtype. **(D)** Heatmap of NMF program scores across all ROIs. Rows correspond to ROIs grouped by histological phenotype and ordered by hierarchical clustering within each group. Left annotation tracks show phenotype, Ki67 labeling index (IHC) and dysplasia grade. The central panel shows scores for each NMF program (red, high; blue, low). Right tracks show semi-quantitative IHC scores (0–3) for MUC2 and CDX2 (intestinal markers) and MUC6 and MUC5AC (gastric markers). The bottom panel shows pairwise Kruskal-Wallis tests of program scores between phenotypes (Intestinal vs Gastric, Intestinal vs PB, Gastric vs Oncocytic, Gastric vs PB, and high-grade Gastric vs PB), with color indicating −log10(p) (*p < 0.05, **p < 0.01, ***p < 0.001). IHC, immunohistochemistry. **(E)** Proportion of *KRAS*-mutant samples and *KRAS* tVAF across histological subtypes. **(F)** Proportion of *GNAS*-mutant samples and *GNAS* tVAF across histological subtypes.

## Results

### Transcriptomic profiling defines two primary molecular axes of IPMN

To capture the full spectrum of PPCN biology, a cohort of 127 patients was assembled, from which 418 microdissected regions were obtained (Fig. 1A). Each microdissected region was composed of a single phenotype and dysplasia grade combination, as defined by the current classification. Depending on the heterogeneity of phenotype and dysplasia, one patient could contribute to one or several regions, providing a comprehensive representation of inter- and intra-tumor IPMN diversity (Fig. 1B). Unsupervised analysis by PCA revealed a clear bipartition of the transcriptomic landscape (Fig. 1C). Intestinal lesions (n=191) were separated from all other lesions along PC1, whereas gastric (n=173), PB (n=43) and oncocytic (n=11) lesions grouped together, sharing transcriptional features (Supplementary Fig. S1A). Dysplasia grade was not associated with either principal component, indicating that it was not a primary axis of transcriptomic variation and supporting the existence of molecular continua beyond conventional histological grading (Fig. 1C and Supplementary Fig. S1B).

To define the main transcriptomic programs of PPCN in an unsupervised manner, non-negative matrix factorization (NMF) was applied to the microdissected profiles, and recurrent programs were identified across multiple runs (Methods). Nine recurrent RNA programs were retained: four present in intestinal lesions, two in gastric and oncocytic lesions, two in gastric and PB lesions, and one broadly expressed mucin-producing program present in all phenotypes (Fig. 1D and Supplementary Table S1). Their distribution recapitulated the bipartition seen by PCA. The four intestinal programs were restricted to intestinal lesions and captured a gradient of transcriptional states, from differentiated to proliferative, whereas gastric, oncocytic and PB lesions broadly shared their programs. The two PB programs were also expressed by high-grade gastric lesions, reducing the transcriptomic separation between these phenotypes in aggressive high-grade lesions. None of the intestinal programs was recapitulated by the pathologically defined dysplasia grade, reinforcing the notion of a continuum rather than a discrete transition during progression.

To link genotype and phenotype, we performed variant calling directly from RNA-seq data, focusing on predefined *KRAS* (G12, G13, G60 and Q61) and *GNAS* (R201) hotspot mutations. Variant allele frequencies reported here are transcriptomic (transcribed variant allele frequencies, tVAFs), reflecting the proportion of mutant reads among expressed transcripts and therefore capturing both mutation clonality and allele-specific expression. *KRAS* and *GNAS* showed distinct mutation patterns. *KRAS* hotspot mutations were detected at similar frequencies across gastric, intestinal, and pancreatobiliary (PB) lesions, with no difference in tVAF between phenotypes (Fig. 1E). In contrast, *GNAS* mutations were enriched in intestinal lesions compared with gastric and PB lesions (intestinal: 137/191 [71.7%] vs gastric/PB: 62/211 [29.4%], p = 2.8e-17; regions with undetermined *GNAS* status excluded). Among mutated regions, *GNAS* transcripts were also more abundant in intestinal regions (mean tVAF = 0.49 vs 0.27, p = 7.3e-10; Fig. 1F), and this abundance scaled with intestinal program scores, most strongly with the canonical intestinal program (r = 0.41, p = 9.7e-10; Supplementary Fig. S1C). These associations are predominant, though not exclusive, since some gastric and PB lesions were also *GNAS*-mutant, while oncocytic lesions were wild-type for both *KRAS* and *GNAS*. Overall, PPCN transcriptomes separate into two broad groups, gastric/PB/oncocytic versus intestinal, a divide reinforced by their preferential mutational drivers, with grade as a secondary feature.

### Morphological motifs mirror and refine the transcriptomic axes

To link transcriptomic programs to tissue morphology, a deep learning model was trained on paired HES images and microdissected RNA sequencing data (Fig. 2A). Although each region of interest (ROI) was selected for morphological uniformity according to current classification criteria, HES images reveal subtle architectural and cytological variations within individual regions that bulk RNA sequencing could not resolve. Tile-level predicted transcriptomics allowed this intra-ROI heterogeneity to be decomposed into ten recurrent morphological motifs, providing a higher-resolution view of the transcriptional landscape within and across lesions (Fig. 2B). The full list of morphological motifs, their description, representative images, and associated RNA program is provided in Supplementary Table S2.

**Figure 2.**
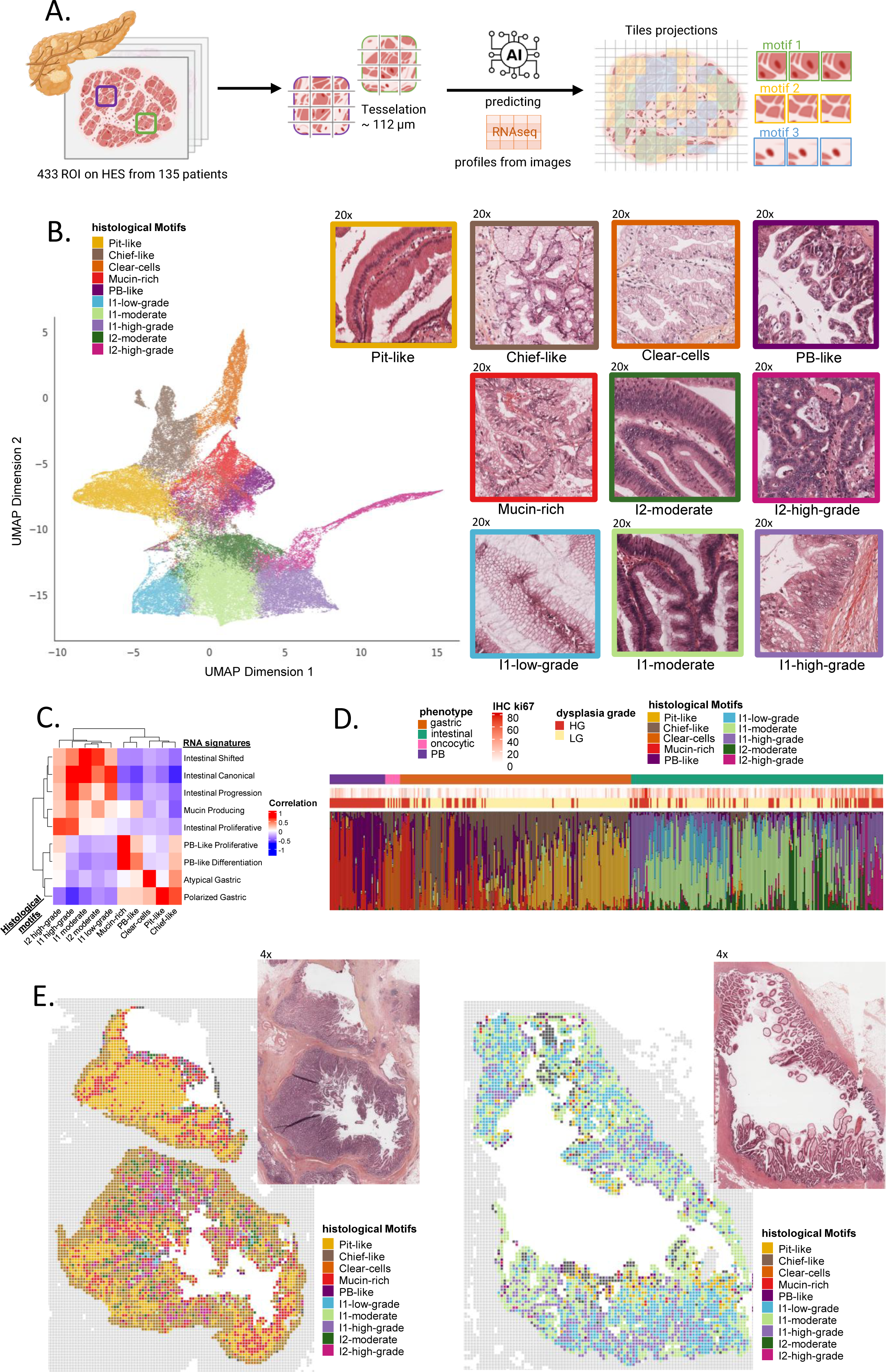
AI-derived morphological motifs recapitulate and extend transcriptomic programs. **(A)** Workflow for morphological motif identification. Digitized FFPE slides were tessellated into tiles, and tile-level RNA expression was predicted using a deep learning model trained on paired HES images and bulk RNA sequencing data. Predicted expression profiles were decomposed into recurrent morphological motifs by consensus NMF. **(B)** UMAP projection of tile-level motif scores based on Euclidean distance, colored by dominant morphological motif. Representative HES images are shown for each motif. **(C)** Correlation matrix between RNA programs and morphological motif scores across all ROIs. **(D)** Heatmap of motif proportion per ROI, ordered by histological subtype and dysplasia grade. **(E)** Spatial projection of morphological motif scores onto representative gastric and intestinal ROIs.

The correspondence between derived transcriptomic programs and morphological motifs established a shared annotation framework for the two modalities (Fig. 2C and Supplementary Table S2). This framework was then used to quantify the spatial composition of tissue regions, revealing the coexistence of multiple transcriptional programs within individual ROIs. Along the gastric axis, the morphological motifs identified in this analysis included *Pit-like* and *Chief-like* motifs, which aligned with the transcriptomic *Polarized Gastric* program. The morphological *Clear-cells* motif aligned with the transcriptomic *Atypical Gastric* program, and *PB-like* and *Mucin-rich* morphological motifs aligned with the two PB transcriptomic programs. Within the intestinal axis, *low-grade* and *high-grade* goblet-rich motifs aligned with the transcriptomic *Intestinal Progression* program, *moderate-grade* motifs with the *Intestinal Shifted* program, and a *highly proliferative* motif with the *Intestinal Proliferative* program. Co-occurrence, heatmap, UMAP and per-phenotype proportion analyses (Fig. 2B, 2D and Supplementary Fig. S1D, S1E), as well as the projection of motifs onto representative gastric and intestinal ROIs (Fig. 2E), consistently organized these motifs into the same two major axes seen in the transcriptomic data. Morphological motifs and RNA programs therefore converge on the same bipartition, separating gastric/PB/oncocytic from intestinal lesions.

### The gastric and pancreatobiliary trajectory recapitulates gastric epithelial polarity and converges toward shared high-grade states

Integrative analysis of ROIs spanning the gastric, pancreatobiliary, and oncocytic phenotypes linked each morphological motif to its transcriptomic program and marker genes (Fig. 3A). Focusing on the gastric phenotype, its low-grade motifs were tested for correspondence to normal stomach epithelial states. Human stomach epithelial cell type signatures (15) were projected onto ROIs classified by their most abundant motif. Each motif was matched to its normal counterpart: *Chief-like* and *Pit-like* dominant ROIs were enriched for the gastric transcriptomic Chief and Pit cell signature, respectively (Fig. 3B). The same result was obtained with mouse reference signatures (16) (Supplementary Fig. S2A). This correspondence also held at the level of tissue architecture. A normal stomach biopsy and a gastric IPMN were profiled by spatial transcriptomics. The same polarized organization was observed in both: *MUC6*-positive, *Chief-like* regions at the gland base and *MUC5AC*-positive, *Pit-like* regions toward the lumen (Fig. 3C). The same polarity was found in two additional low-grade gastric IPMNs profiled by spatial transcriptomics (Supplementary Fig. S2B; summarized in Fig. 3D). The two motifs were consistently identified as distinct states, but their relative proportions varied across lesions, forming a continuum from *Pit-like* to *Chief-like* compositions. This resemblance extended beyond the two mucins used to define the motifs. Across the cohort, canonical markers of stomach polarity were themselves polarized within gastric IPMN: foveolar markers *(MUC5AC, TFF1, NKX6-2)* were more highly expressed in *Pit-like* dominant ROIs, and gland-base markers *(MUC6, TFF2, HOMER2)* in *Chief-like* dominant ROIs (Supplementary Fig. S2C and Supplementary Table S3). Genotype, in contrast, was not evenly distributed along the continuum: *GNAS* mutations were more frequent in *Pit-like* dominant than in *Chief-like* dominant ROIs (p = 0.005) and carried a higher mutant transcript fraction (p = 7.7 × 10⁻⁶; Supplementary Fig. S2D), whereas both groups remained predominantly *KRAS*-mutant (Fig. 3A). Low-grade gastric IPMN therefore closely resembles normal stomach epithelium, while showing variable *Pit-like/Chief-like* compositions that may be driven by GNAS activity.

**Figure 3.**
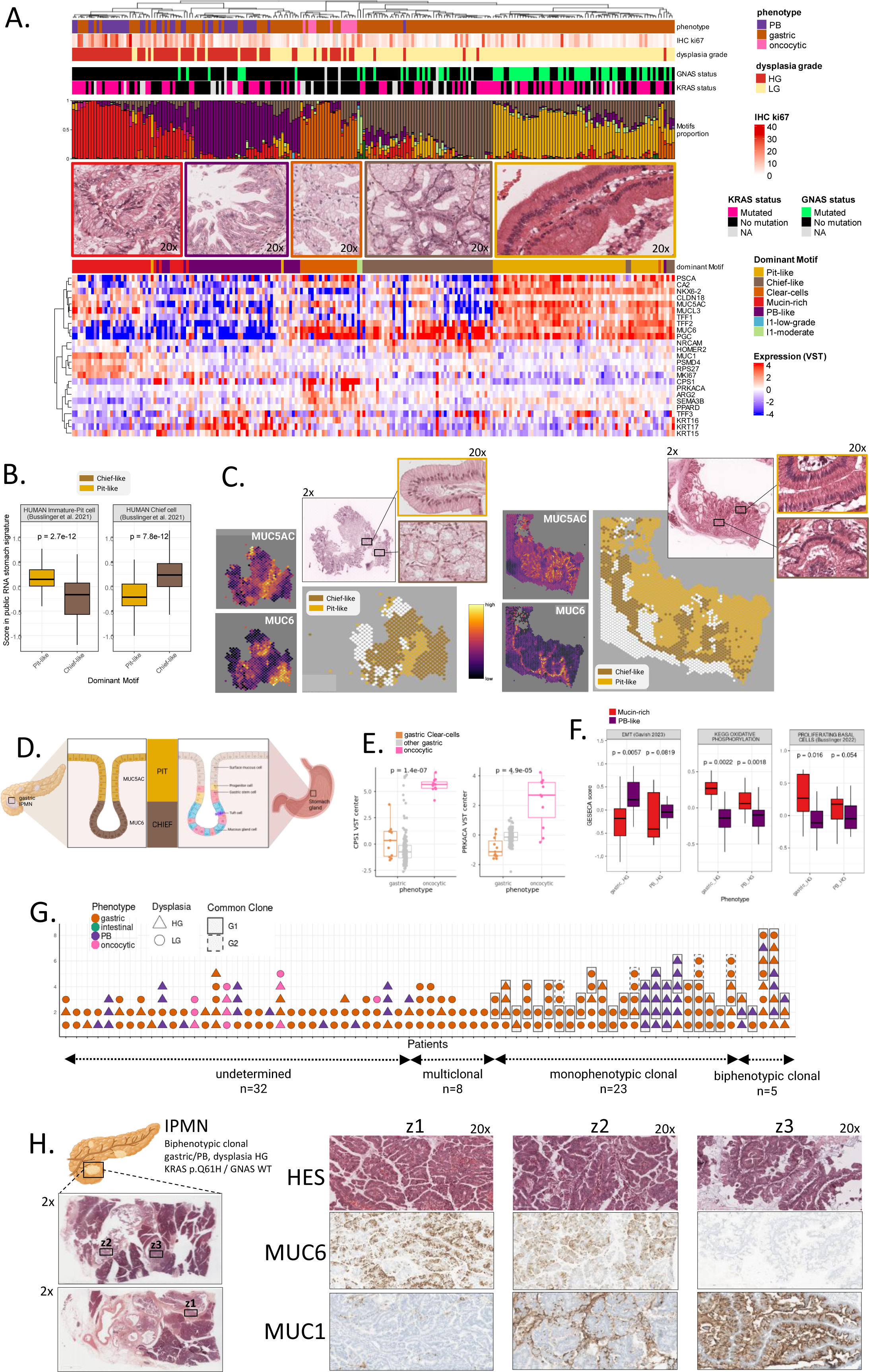
The gastric/PB axis recapitulates gastric epithelial polarity and reveals distinct biological trajectories. **(A)** Integrative heatmap of ROIs spanning the gastric, pancreatobiliary (PB) and oncocytic phenotypes. Columns correspond to ROIs ordered by hierarchical clustering. From top to bottom: phenotype, Ki67 labeling index (IHC) and dysplasia grade; *GNAS* and *KRAS* mutation status; motif proportions per ROI (stacked bars); representative HES images of each dominant motif, framed in the corresponding motif color; dominant motif; and expression of representative marker genes (VST-normalized, genes ordered by hierarchical clustering). **(B)** GESECA enrichment analysis of human gastric pit and chief cell reference signatures in *Pit-like* and *Chief-like* dominant IPMNs. **(C)** Spatial projection of the *Pit-like* and *Chief-like* motifs onto spatial transcriptomics data from a normal gastric biopsy and a gastric IPMN. For each section: spatial expression of *MUC5AC* and *MUC6*; HES overview with magnified insets of *Pit-like* (yellow frame) and *Chief-like* (brown frame) regions; and spatial distribution of the NSF factors most correlated with the *Pit-like* (yellow) and *Chief-like* (brown) motifs (IPMN spots below both thresholds in white; non-IPMN spots in grey). **(D)** Schematic representation of epithelial polarity in normal gastric glands and gastric IPMN, showing the spatial organization of the *Chief-like* and *Pit-like* programs. **(E)** Expression of the oncocytic markers *CPS1* and *PRKACA* in oncocytic IPMNs, gastric IPMNs enriched for the *Clear-cells* motif, and other gastric IPMNs. **(F)** GESECA enrichment analysis of *Mucin-rich* and *PB-like* dominant samples. **(G)** Multi-region clonality analysis. Each column corresponds to a patient and each symbol to an ROI, colored by histological phenotype and shaped by dysplasia grade (triangle, HG; circle, LG). ROIs sharing identical *KRAS* and/or *GNAS* mutations are boxed as a common clone (solid line, first clone, G1; dashed line, second clone within the same patient, G2). Patients were classified as undetermined (n = 32; clonal relationship could not be inferred because of insufficient sequencing depth or driver tVAF below the detection threshold), multiclonal (n = 8; ROIs carrying distinct driver mutations), monophenotypic monoclonal (n = 23; clonally related ROIs of a single phenotype) or biphenotypic monoclonal (n = 5; clonally related gastric and PB ROIs). **(H)** Immunohistochemical staining for MUC6 and MUC1 in three clonally related lesions from the same biphenotypic patient, spanning low-grade gastric, high-grade gastric and PB phenotypes.

A subset of gastric lesions shared transcriptomic features with oncocytic cases, through their common contribution to the *Atypical Gastric* program and the *Clear-cells* motif (Fig. 3A and Supplementary Fig. S3A). This overlap is likely due to common metabolic states: the *Clear-cells* motif was enriched for the urea cycle and glycolysis (Supplementary Fig. S3B), consistent with the mitochondrion-rich biology of oncocytic epithelium. This overlap was restricted to transcriptional features and did not extend to the molecular alterations and markers characteristic of oncocytic lesions. *PRKACA or PRKACB* rearrangements were detected in all 11 oncocytic lesions and in none of the 11 *Clear-cells* dominant gastric ROIs. The oncocytic markers *CPS1* (Hepar1) and *PRKACA* were also expressed at significantly higher levels in oncocytic than in gastric lesions, whether *Clear-cells* dominant or not (Fig. 3E). *Clear-cells* dominant gastric lesions are therefore gastric IPMN, not oncocytic lesions: they share an oncocytic-like metabolic signature but lack the fusion transcripts and markers that define oncocytic identity.

High-grade gastric and PB lesions were characterized by the same two morphological motifs, *Mucin-rich* and *PB-like*, present in both phenotypes in comparable proportions (Fig. 3A and Supplementary Fig. S3C). The *Mucin-rich* motif was characterized by high *MUC1* expression and enrichment for metabolic reprogramming and proliferative cell signature (15) (Fig. 3F and Supplementary Fig. S3D, Supplementary Table S4). The *PB-like* motif showed increased expression of *KRT15, KRT16* and *KRT17* and enrichment for an epithelial-to-mesenchymal transition signature (17) (Fig. 3F and Supplementary Fig. S3D, Supplementary Table S4). In contrast to the basal Chief-like and apical Pit-like organization of low-grade gastric lesions, neither high-grade motif was polarized along the epithelium, and MUC1 staining extended along the entire papilla. The shared morphological motifs across high-grade gastric and PB lesions raised the possibility that PB lesions could arise from gastric lesions within a common clone. We therefore screened multi-region samples within the same patients for shared *KRAS* and *GNAS* mutations (Fig. 3G). Clonality-resolvable events were found in 36 of 68 patients, and among them 5 had clonal events spanning different phenotypes (biphenotypic monoclonal), with shared clonal *KRAS* or *GNAS* mutations across gastric and PB ROIs. Thus, gastric and PB phenotypes can share a common clonal origin, suggesting that PB lesions may derive from gastric lesions through phenotypic evolution. This relationship was further illustrated at the protein level in one biphenotypic monoclonal case comprising three clonally related lesions spanning low-grade gastric, high-grade gastric and PB phenotypes (Fig. 3H). MUC6 and MUC1 showed opposing distributions across these lesions, with strong MUC6 staining in the gastric lesions and progressively greater MUC1 staining toward the PB lesion. Together, these findings support a possible phenotypic progression from low-grade gastric to high-grade gastric and PB states within a common clonal lineage.

Integrative analysis of ROIs spanning the gastric, pancreatobiliary, and oncocytic phenotypes linked each morphological motif to its transcriptomic program and marker genes (Fig. 3A). Focusing on the gastric phenotype, its low-grade motifs were tested for correspondence to normal stomach epithelial states. Human stomach epithelial cell type signatures (15) were projected onto ROIs classified by their most abundant motif. Each motif was matched to its normal counterpart: *Chief-like* and *Pit-like* dominant ROIs were enriched for the gastric transcriptomic Chief and Pit cell signature, respectively (Fig. 3B). The same result was obtained with mouse reference signatures (16) (Supplementary Fig. S2A). This correspondence also held at the level of tissue architecture. A normal stomach biopsy and a gastric IPMN were profiled by spatial transcriptomics. The same polarized organization was observed in both: *MUC6*-positive, *Chief-like* regions at the gland base and *MUC5AC*-positive, *Pit-like* regions toward the lumen (Fig. 3C). The same polarity was found in two additional low-grade gastric IPMNs profiled by spatial transcriptomics (Supplementary Fig. S2B; summarized in Fig. 3D). The two motifs were consistently identified as distinct states, but their relative proportions varied across lesions, forming a continuum from *Pit-like* to *Chief-like* compositions. This resemblance extended beyond the two mucins used to define the motifs. Across the cohort, canonical markers of stomach polarity were themselves polarized within gastric IPMN: foveolar markers *(MUC5AC, TFF1, NKX6-2)* were more highly expressed in *Pit-like* dominant ROIs, and gland-base markers *(MUC6, TFF2, HOMER2)* in *Chief-like* dominant ROIs (Supplementary Fig. S2C and Supplementary Table S3). Genotype, in contrast, was not evenly distributed along the continuum: *GNAS* mutations were more frequent in *Pit-like* dominant than in *Chief-like* dominant ROIs (p = 0.005) and carried a higher mutant transcript fraction (p = 7.7 × 10⁻⁶; Supplementary Fig. S2D), whereas both groups remained predominantly *KRAS*-mutant (Fig. 3A). Low-grade gastric IPMN therefore closely resembles normal stomach epithelium, while showing variable *Pit-like/Chief-like* compositions that may be driven by GNAS activity.

A subset of gastric lesions shared transcriptomic features with oncocytic cases, through their common contribution to the *Atypical Gastric* program and the *Clear-cells* motif (Fig. 3A and Supplementary Fig. S3A). This overlap is likely due to common metabolic states: the *Clear-cells* motif was enriched for the urea cycle and glycolysis (Supplementary Fig. S3B), consistent with the mitochondrion-rich biology of oncocytic epithelium. This overlap was restricted to transcriptional features and did not extend to the molecular alterations and markers characteristic of oncocytic lesions. *PRKACA or PRKACB* rearrangements were detected in all 11 oncocytic lesions and in none of the 11 *Clear-cells* dominant gastric ROIs. The oncocytic markers *CPS1* (Hepar1) and *PRKACA* were also expressed at significantly higher levels in oncocytic than in gastric lesions, whether *Clear-cells* dominant or not (Fig. 3E). *Clear-cells* dominant gastric lesions are therefore gastric IPMN, not oncocytic lesions: they share an oncocytic-like metabolic signature but lack the fusion transcripts and markers that define oncocytic identity.

High-grade gastric and PB lesions were characterized by the same two morphological motifs, *Mucin-rich* and *PB-like*, present in both phenotypes in comparable proportions (Fig. 3A and Supplementary Fig. S3C). The *Mucin-rich* motif was characterized by high *MUC1* expression and enrichment for metabolic reprogramming and proliferative cell signature (15) (Fig. 3F and Supplementary Fig. S3D, Supplementary Table S4). The *PB-like* motif showed increased expression of *KRT15, KRT16* and *KRT17* and enrichment for an epithelial-to-mesenchymal transition signature (17) (Fig. 3F and Supplementary Fig. S3D, Supplementary Table S4). In contrast to the basal Chief-like and apical Pit-like organization of low-grade gastric lesions, neither high-grade motif was polarized along the epithelium, and MUC1 staining extended along the entire papilla. The shared morphological motifs across high-grade gastric and PB lesions raised the possibility that PB lesions could arise from gastric lesions within a common clone. We therefore screened multi-region samples within the same patients for shared *KRAS* and *GNAS* mutations (Fig. 3G). Clonality-resolvable events were found in 36 of 68 patients, and among them 5 had clonal events spanning different phenotypes (biphenotypic monoclonal), with shared clonal *KRAS* or *GNAS* mutations across gastric and PB ROIs. Thus, gastric and PB phenotypes can share a common clonal origin, suggesting that PB lesions may derive from gastric lesions through phenotypic evolution. This relationship was further illustrated at the protein level in one biphenotypic monoclonal case comprising three clonally related lesions spanning low-grade gastric, high-grade gastric and PB phenotypes (Fig. 3H). MUC6 and MUC1 showed opposing distributions across these lesions, with strong MUC6 staining in the gastric lesions and progressively greater MUC1 staining toward the PB lesion. Together, these findings support a possible phenotypic progression from low-grade gastric to high-grade gastric and PB states within a common clonal lineage.

### The intestinal trajectory follows two evolutionary routes with distinct origins

All intestinal lesions shared a specific transcriptomic program, *Intestinal Canonical*, with uniform expression of *MUC2*, *CDX2* and *SPINK4* (Fig. 1D, 4A). This program distinguished intestinal lesions from the other PPCN phenotypes, but did not distinguish them from each other. Two additional transcriptomic programs, *Intestinal Progression* and *Intestinal Shifted*, defined two distinct states along the intestinal axis (Fig. 4A, 4B and Supplementary Fig. S4A). *Progression* lesions showed an almost exclusively intestinal phenotype, with intestinal programs and intestinal morphology (Fig. 4A, 4C). In contrast, *Shifted* lesions showed a biphenotypic phenotype. They combined intestinal and gastric transcriptomic programs and retained *Chief-like* and *PB-like* morphological motifs (Fig. 4A, 4C). They were also enriched for metabolic rather than proliferative programs (Supplementary Fig. S4B and Supplementary Table S5). The two groups also differed genetically and clinically. *KRAS* mutations were found in 30 of 65 *Shifted* lesions (46%) compared with 4 of 32 *Progression* lesions (13%, p = 0.0013) (Fig. 4D). *Shifted* lesions were also more frequently found in patients with both gastric and intestinal histologies: 20 of 65 (31%) compared with 2 of 32 (6%, p = 0.006) (Fig. 4D). Together, these findings link the *Shifted* state to gastric features and they raise the possibility that *Intestinal Shifted* lesions may arise from a lesion with a previous gastric phenotype.

**Figure 4.**
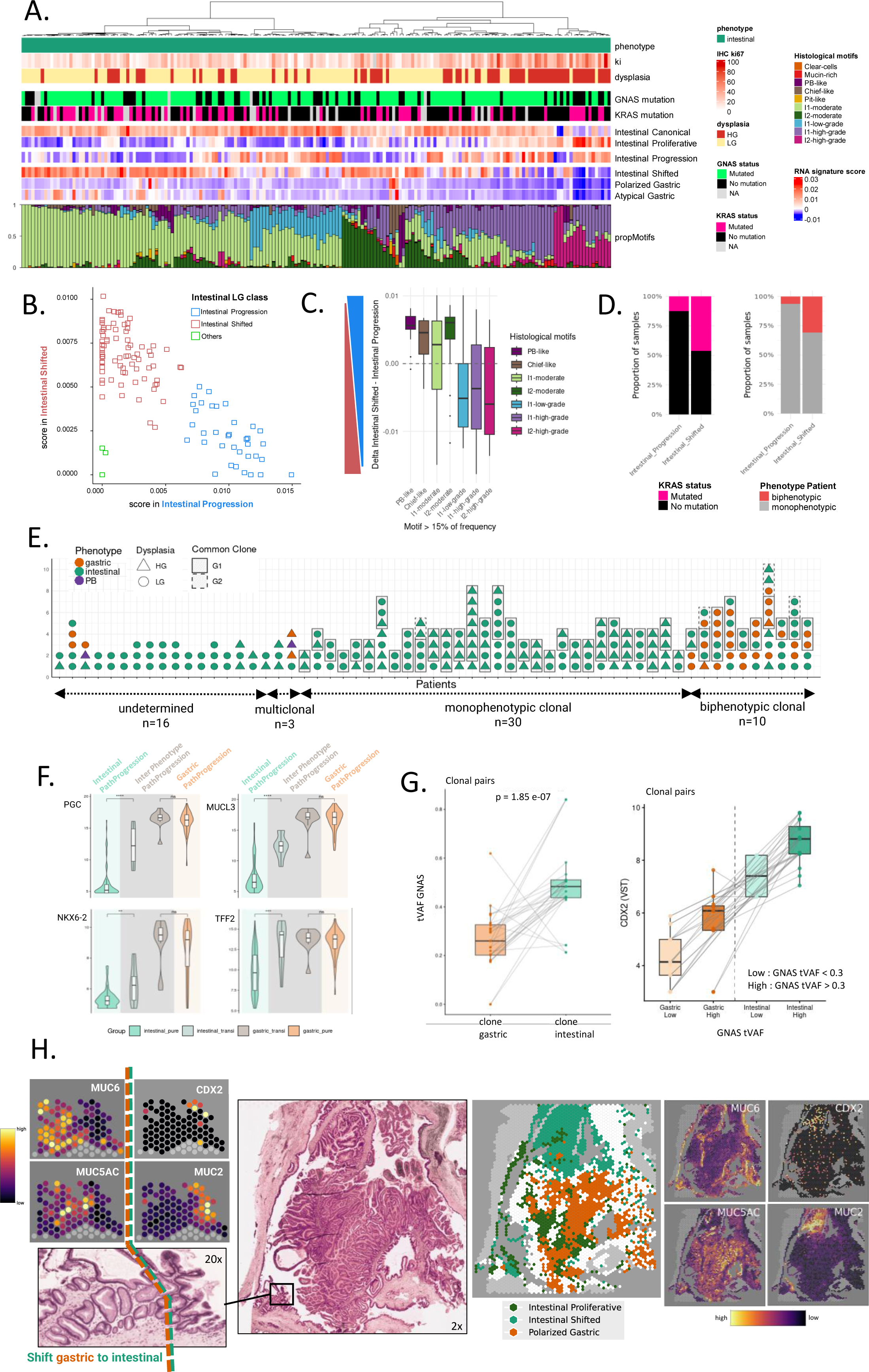
Distinct evolutionary trajectories shape intestinal IPMN progression. **(A)** Integrative heatmap of intestinal ROIs. Columns correspond to ROIs ordered by hierarchical clustering. From top to bottom: phenotype, Ki67 labeling index (IHC) and dysplasia grade; *GNAS* and *KRAS* mutation status; RNA program scores (*Intestinal Canonical, Intestinal Proliferative, Intestinal Progression, Intestinal Shifted, Polarized Gastric* and *Atypical Gastric*); and histological motif proportions per ROI (stacked bars). **(B)** Model-based clustering (Mclust) of low-grade intestinal ROIs based on *Intestinal Progression* and *Intestinal Shifted* program scores. ROIs not assigned to either cluster are shown as Others **(C)** Difference between *Intestinal Shifted* and *Intestinal Progression* program scores in intestinal ROIs, grouped by histological motifs present at > 15% frequency. Positive values indicate a predominance of the *Intestinal Shifted* program and negative values a predominance of the *Intestinal Progression* program. **(D)** *KRAS* mutation frequency and proportion of ROIs from mixed gastric/intestinal patients in the *Intestinal Progression* and *Intestinal Shifted* groups. **(E)** Multi-region clonality analysis of patients with intestinal lesions. Each column corresponds to a patient and each symbol to an ROI, colored by histological phenotype and shaped by dysplasia grade (triangle, HG; circle, LG). ROIs sharing identical *KRAS* and/or *GNAS* mutations are boxed as a common clone (solid line, G1; dashed line, second clone within the same patient, G2). Patients were classified as undetermined (n = 16; clonal relationship could not be inferred because of insufficient sequencing depth or driver tVAF below the detection threshold), multiclonal (n = 3), monophenotypic monoclonal (n = 30; intestinal-restricted clones) or biphenotypic monoclonal (n = 10; clonally related gastric and intestinal ROIs). **(F)** Expression of gastric lineage markers (*PGC, MUCL3, NKX6-2, TFF2*) in four groups of ROIs: intestinal ROIs without a clonally related gastric lesion (intestinal path), intestinal and gastric ROIs belonging to clonally related gastric-intestinal pairs (inter-phenotype path), and gastric ROIs without a clonally related intestinal lesion (gastric path). Clonal relationships were defined by shared *KRAS* and/or *GNAS* mutations (Wilcoxon rank-sum test; *p < 0.05, **p < 0.01, ***p < 0.001, ****p < 0.0001; ns, not significant). **(G)** Clonally related gastric–intestinal lesion pairs. Left: *GNAS* tVAF in the gastric and intestinal ROIs of each pair (lines connect paired ROIs; Wilcoxon signed-rank test). Right: *CDX2* expression (VST) in the same ROIs, split into four groups by phenotype and *GNAS* tVAF (gastric low, gastric high, intestinal low, intestinal high; low, tVAF < 0.3; high, tVAF ≥ 0.3). **(H)** Spatial transcriptomic analysis of a clonally validated mixed gastric–intestinal IPMN. HES overview (2×) with a magnified view (20×) of the gastric-to-intestinal transition zone (dashed line), and spatial expression of *MUC6, MUC5AC, CDX2* and *MUC2* in this region. Right: spatial distribution of the *Polarized Gastric, Intestinal Shifted* and *Intestinal Proliferative* programs across the section (IPMN spots below all thresholds in white; non-IPMN spots in grey), and corresponding whole-section expression of *MUC6, MUC5AC, CDX2* and *MUC2*.

Clonal relationships were inferred from shared *KRAS* and *GNAS* variants across multi-region samples (Fig. 4E). Clonality-resolvable events were found in 43 of 59 patients with intestinal lesions. Among them, 10 had clonal events spanning different phenotypes (biphenotypic monoclonal), with gastric and intestinal lesions sharing a clonal *KRAS* and/or *GNAS* mutation. Intestinal and PB lesions were never found to be clonally related across the 43 informative patients, suggesting that this absence is unlikely to be explained by detection limits alone. In these gastric-intestinal biphenotypic events, the intestinal lesions retained molecular features of their gastric counterparts, expressing higher *PGC, MUCL3, TFF2* and *NKX6-2*, together with a higher *Pit-like* contribution (Fig. 4A, 4F). Clonally related gastric and intestinal lesions were also closer to each other in PCA space (Supplementary Fig. S4C).

Symmetrically, gastric lesions sharing an intestinal clone expressed higher levels of intestinal genes compared to pure, unshifted, gastric lesions (Supplementary Fig. S4D). Within these pairs, *GNAS* tVAF was higher in intestinal lesions than in gastric lesions, and CDX2 expression increased with *GNAS* tVAF within each phenotype, peaking in intestinal lesions with high *GNAS* tVAF (Fig. 4G). This is consistent with the association of *GNAS* mutations with intestinal differentiation rather than with dysplasia grade (9). These associations point to a link between gastric and intestinal lesions, consistent with the biphenotypic profile of *Shifted* lesions.

These relationships were then examined in tissue by spatial transcriptomics, on clonally validated biphenotypic gastric-intestinal IPMN (Fig. 4H). Within a single lesion, low-grade *Polarized Gastric* regions were adjacent to *Intestinal Shifted* territory, which in turn bordered high-grade areas expressing the *Intestinal Proliferative* program. At higher magnification, two of these states met within one continuous epithelium: along a single papilla, the epithelial sheet transitioned from *MUC6*-positive *Chief-like* to *MUC5AC*-positive *Pit-like* territory and then, without interruption, to a *MUC2*-positive *Intestinal Shifted* segment. The shifted state was therefore not simply adjacent to a gastric lesion but continuous with the gastric epithelium. This transition was independently supported by immunohistochemistry in a monoclonal biphenotypic IPMN, in which MUC6 and CDX2 showed reciprocal patterns across the gastric-intestinal interface (Supplementary Fig. S4E, S4F): MUC6 was predominantly retained in the gastric compartment, CDX2 was enriched in the intestinal compartment, and the transition zone was marked by loss of MUC6 and acquisition of CDX2. Together, these clonal, spatial and marker data support two routes to the *Intestinal Canonical* state: a direct route, and an *Intestinal Shifted* route that retains molecular, morphological, genetic and spatial traces of a gastric past.

### IPMN trajectories map onto PDAC subtypes and are detectable in mucus

To investigate potential malignant endpoints of IPMN trajectories, IPMN programs and public PDAC signatures (19, 22–26) were projected onto independent PDAC cohorts (TCGA PAAD (18), CPTAC (21), Maurer (20), and Puleo (19)) (Fig. 5A). Associations emerged between IPMN trajectories and PDAC molecular states. Gastric-associated programs, including *Polarized Gastric*, *Atypical Gastric*, and *PB-like Differentiation*, preferentially aligned with Classical A PDAC signatures, consistent with their differentiated glandular features. Intestinal programs aligned more with Classical B, the least represented subtype among metastatic tumors (25), with *Intestinal Progression, Intestinal Canonical,* and *Intestinal Shifted* all following this trend. The *Intestinal Shifted* program displayed an intermediate profile, showing contribution from both Classical A and Classical B signatures, consistent with its mixed gastric and intestinal identity. Proliferative programs, including *PB-like Proliferative* and *Intestinal Proliferative*, showed the strongest association with Basal PDAC subtypes. The *Mucin Producing* program showed preferential similarity with Classical A rather than Basal states (Fig. 5A). These relationships are summarized in an integrated model of IPMN evolutionary trajectories and their putative PDAC endpoints (Fig. 5B).

**Figure 5.**
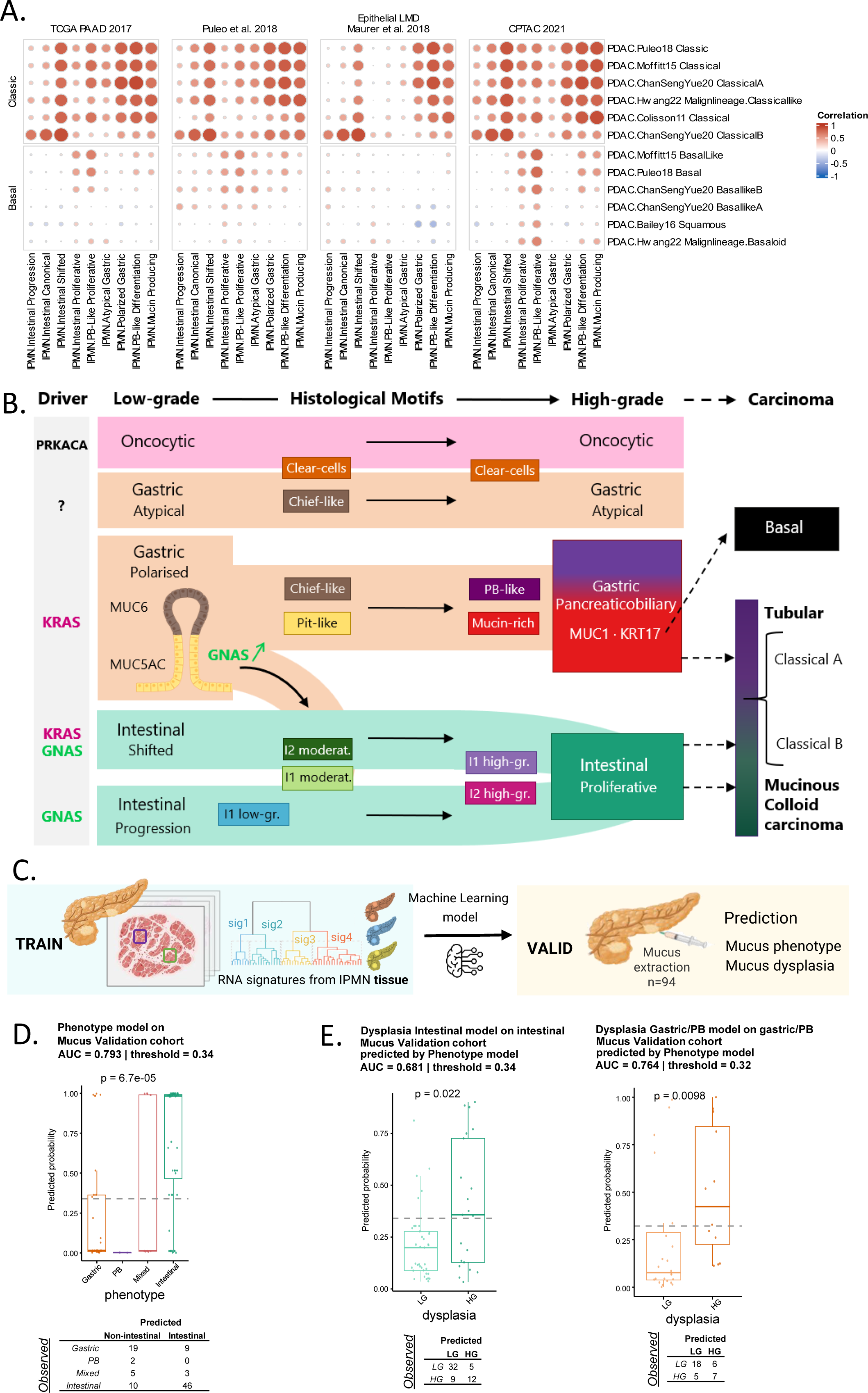
IPMN trajectories link to PDAC subtypes and enable mucus-based prediction. **(A)** Projection of IPMN transcriptomic programs onto TCGA (18), Puleo (19), Maurer (20), and CPTAC (21) PDAC cohorts. **(B)** Integrated model of IPMN evolutionary trajectories linking gastric, oncocytic, pancreatobiliary, *Intestinal Shifted*, and *Intestinal Progression* routes to downstream cancer states. **(C)** Workflow of mucus prediction model trained on tissue RNA programs and validated on independent mucus samples annotated for phenotype and dysplasia. **(D)** Phenotype prediction scores in the mucus validation cohort, by observed phenotype. **(E)** Dysplasia prediction scores in the mucus validation cohort, from the intestinal model in samples predicted as intestinal (left) and from the non-intestinal (gastric/pancreatobiliary) model in samples predicted as non-intestinal (right). In (D-E), dashed lines indicate classification thresholds defined on tissue cross-validation (Youden index) and applied unchanged to the mucus cohort. Contingency tables show observed versus predicted classes. p-values: Kruskal-Wallis (D) and Wilcoxon rank-sum (E). AUC, area under the ROC curve; LG, low-grade; HG, high-grade dysplasia.

To assess the translational potential of these phenotype-specific paths, tissue-derived RNA programs were used to train a two-step classifier on tissue samples, which was then applied to RNA extracted from IPMN mucus (Fig. 5C). The first step assigned a phenotype (intestinal versus non-intestinal), and the second applied a dedicated dysplasia model within each predicted phenotype. The classifier was applied to an independent cohort of 94 IPMN mucus samples profiled by RNA sequencing (56 intestinal, 30 gastric or PB, 8 biphenotypic). Phenotype prediction scores differed between phenotypes (p = 6.7 × 10^-5^), and within each branch the tissue-based dysplasia models differentiated low-from high-grade lesions (intestinal p = 0.022; gastric/PB, p = 0.0098) (Fig. 5D, 5E). By comparison, a unified dysplasia model trained without phenotype assignment reached lower performance in mucus (AUC 0.68-0.70) and did not significantly separate low-from high-grade gastric/PB or biphenotypic lesions (Supplementary Fig. S5A). IPMN transcriptional programs defined in tissue are therefore detectable in mucus-derived RNA and retain phenotype- and grade-associated information, supporting mucus-based profiling as a minimally invasive approach for risk stratification pending validation in larger cohorts.

## Discussion

PPCN are managed under a linear model, in which low-grade dysplasia progresses to high-grade dysplasia and eventually to invasive carcinoma. This model underlies current surgical criteria, yet molecular studies have consistently failed to identify a signature separating low from high grade (7, 8, 13). Our data support a different model: rather than a single sequence, PPCN comprises distinct trajectories that can diverge toward different high-grade endpoints, most trajectories arising from a shared low-grade gastric initial state. Grade is thereby a shared label applied to biologically different progression routes, which explains why it has resisted molecular definition and points to a practical alternative: resolve phenotype first, then grade within it.

The proposed model relies first on the nature of the starting point. Low-grade gastric IPMN preserves the multicellular luminal-to-basal polarity of the normal stomach gland, with *Pit-like* and *Chief-like* motifs mirroring foveolar and chief cell identities. Deep learning-based morphology showed that this polarity was lost with progression, as high-grade lesions became dominated by the non-polarized *Mucin-rich* and *PB-like motifs*. From this normal-mimicking ground state, progression diverged along a limited number of routes, which we define by their terminal phenotype.

The first route remained within the gastric trajectory. Along the gastric/PB progression trajectory, gastric lesions lost their gland-like organization and converged toward a high-grade state shared with PB lesions, so that high-grade gastric and PB lesions were largely intermixed along the major molecular axes. This route therefore progresses to high grade without changing lineage, which is consistent with the recent spatial transcriptomic study rooting a continuum toward PB phenotypes in a heterogeneous gastric state (27).

The second route involved a change of lineage, and *GNAS* appears to be its main determinant. *GNAS* mutations were enriched in lesions with a dominant *Pit-like* compartment, and because *GNAS* and cAMP signaling drive intestinal differentiation (28, 29), their activation may provide a permissive context for *CDX2*-mediated reprogramming (30) and entry into a shifted route toward the intestinal phenotype. Several independent observations support such a gastric-to-intestinal shift: biphenotypic lesions with identical driver mutations (*KRAS, GNAS*), retention of gastric transcriptional traces in shifted lesions, and an increase in mutated *GNAS* allele transcription along the transition. Spatial transcriptomics further showed that shifted intestinal and gastric cells coexisted within the same continuous epithelium, supporting an *in-situ* conversion rather than the collision of two independent lesions.

Importantly, these *shifted intestinal* lesions are not equivalent to intestinal lesions arising without detectable gastric ancestry. Both converged on the same *Intestinal Proliferative* program. However, “pure” intestinal lesions projected mostly onto Classical B PDAC, whose prototypic form is intestinal colloid carcinoma, whereas the shifted signature projected equally onto Classical A and B, the former being defined by gastric epithelial markers such as *TFF1* and *CLDN18*. The intestinal IPMN phenotype is thus reached by two distinct routes rather than one. Which cell initiates the switch, and under which conditions, remains open; candidate hypotheses include a distinct cell of origin or interactions with a specific microenvironment. These questions cannot be settled from resection specimens, where the transition is only observed after the fact, and will require lineage tracing in models that reproduce PPCN biology.

Overall, clonality analysis supports the different trajectories. Clonally related lesions repeatedly linked gastric with PB states and gastric with intestinal states, consistent with the transcriptomic proximity of high-grade gastric and PB lesions and with the gastric traces retained in shifted intestinal lesions. Conversely, intestinal and PB lesions were essentially never clonally related, and this separation held at the transcriptomic and morphological levels: PB and intestinal lesions shared no RNA program or motif and were separated along PC1 (Fig. 1C, 1D and 2D). Shared driver mutations and transcriptomic evidence thus converge on a separation between the two trajectories downstream of the gastric ground state. This phenotypic view complements multi-region whole-genome sequencing studies of IPMN clonal evolution (31). Whether comparable routes organize other pancreatic precursors, such as pancreatic intraepithelial neoplasia (PanIN), which shares features with low-grade gastric IPMN (32), remains to be explored.

IOPN extends this framework to a third, genetically distinct entity. Although separated from IPMN in the WHO classification, they share clinical and radiological presentation with PPCN, may progress to PDAC, and should therefore be managed within the same overall framework. Their transcriptomic profiles overlapped with *Clear-cells* dominant gastric lesions, indicating biological similarity, yet their path is distinct: *PRKACA/PRKACB* fusions were confined to IOPN and absent from every gastric lesion screened. Because IOPN lacks the *KRAS* and *GNAS* mutations on which our clonal inference rests, whether they share an ancestor with gastric precursors cannot be tested with a variant-based approach.

The identified trajectories may extend beyond the precursor stage. Some transcriptional programs associated with molecular subtypes of invasive PDAC were already detectable at the neoplastic stage, indicating that clinically relevant features of invasive disease may emerge before invasion. Bulk PDAC transcriptomes are influenced by tumor purity and stromal composition, but the reproducibility of these associations across four independent cohorts argues that they reflect underlying lineage relationships rather than tumor composition alone. Establishing whether a given carcinoma descends from a specific precursor trajectory will require cohorts in which genetically related precursor and invasive components are profiled in the same patient.

The clinical value of this framework ultimately depends on whether trajectories can be identified before surgery. In our data, grade alone provided limited separation along major molecular axes, whereas phenotype and trajectory showed stronger associations. Once phenotype was resolved, grade became more informative: phenotype-specific models separated low-from high-grade lesions in both branches whereas a single model trained on all lesions performed less well in mucus (Supplementary Fig. S5A). As a proof of concept, we showed that this information could be retrieved from mucus aspirated from resected specimens. Translating this into preoperative practice will require a prospective study collecting pancreatic juice or cyst mucus before surgery, for instance after secretin stimulation (33), together with the design and validation of a dedicated predictive model. Mixed or ambiguous phenotypes in mucus may limit this approach without trajectory assignment. Nevertheless, this framework shifts the clinical question from how advanced a lesion appears to which biological trajectory it is following, and could help identify lesions at higher risk of progression without relying on grade alone.

## Methods

### Patient samples and histological review

This study was approved by the Institutional Review Board of the Paris Nord Val-de-Seine University Hospitals Group and Université Paris Cité (IRB 2024-A02816-41). We retrieved IPMN resections performed between 2013 and 2023 at Beaujon Hospital (Clichy, France). Two expert pancreatic pathologists (JC, AC) reviewed all slides and assigned histological subtype and dysplasia grade according to the 6th WHO classification (Supplementary Methods). Regions of interest (ROIs) of homogeneous subtype and grade, free of invasive carcinoma, were selected for microdissection (Fig. 1A), and subtype was confirmed by immunohistochemistry (IHC) for MUC2, MUC5AC, MUC6 and CDX2.

### Histological Staining, Immunohistochemistry, and Digitization

HES and IHC staining were performed on automated platforms (Leica ST5020 and Roche BenchMark ULTRA), and slides were digitized on an Aperio AT Turbo scanner (Leica). MUC2, MUC5AC, MUC6, CDX2, MUC1 and CPS1 staining was scored semi-quantitatively (0–3), and the Ki67 labeling index was measured with QuPath v0.4.

### Tissue processing, RNA extraction, and sequencing

ROIs were microdissected from 10-µm FFPE sections with the Avenio Millisect system (Roche Diagnostics). RNA was extracted with the Quick-RNA FFPE Kit (Zymo Research), and libraries were built with the SMARTer Stranded Total RNA-Seq Kit Pico Input Mammalian (Takara Bio) and sequenced on a NovaSeq X (Illumina). Reads were aligned to GRCh38 (Ensembl 107) with STAR and summarized with featureCounts; counts were normalized by variance-stabilizing transformation (VST; DESeq2), and PCA used the 1,000 most variable genes. Mucus samples from the validation cohort were collected and processed as described in the Supplementary Methods.

### Fusion detection

All 11 oncocytic lesions and all 11 gastric lesions dominated by the *Clear-cells* motif were screened for *PRKACA* and *PRKACB* rearrangements with the Archer FusionPlex Pan Solid Tumor v2 panel (Integrated DNA Technologies).

### Sample selection and quality control

Of 433 ROIs microdissected from 135 patients, 15 with fewer than 20,000 detected genes were excluded. The 418 remaining ROIs from 127 patients included 191 intestinal (71 high-grade), 173 gastric (36 high-grade), 43 pancreatobiliary (all high-grade, as defined by the WHO classification) and 11 oncocytic (5 high-grade) regions.

### RNA-Based Variant Calling

*KRAS* and *GNAS* hotspot mutations were called directly from RNA-seq data with RNApipeline v1.7 (https://github.com/genehetx/RNApipeline), which computes read depth and tVAF at predefined hotspot positions with SAMtools. A mutation was called when supported by at least 10 reads and a tVAF of at least 0.2.

### Clonality Inference

For each pair of ROIs from the same patient, *KRAS* and *GNAS* hotspot mutations were compared at the nucleotide level. Pairs sharing at least one identical mutation were considered clonal, pairs carrying only different mutations non-clonal, and pairs lacking informative mutations undetermined. Patients were then classified as monoclonal (monophenotypic or biphenotypic), multiclonal or undetermined (Supplementary Methods).

### Differential Expression and Pathway Analysis

Differential expression was tested with DESeq2. Gene set enrichment was assessed with GSEA on ranked gene lists and with GESECA (fgsea) on the normalized expression matrix; mouse reference signatures (16) were mapped to human orthologs.

### Statistical analysis

All tests were two-sided, with p < 0.05 considered significant and Benjamini-Hochberg correction applied where indicated. The test used for each analysis is given in the figure legends and Supplementary Methods.

### Identification of Recurrent RNA Programs by Non-Negative Matrix Factorization

Recurrent RNA programs were identified by NMF of the expression matrix, run with k = 2 to 30 and random initialization. Programs from all runs were clustered by Jaccard distance, with the number of clusters chosen by silhouette width, and one representative program was retained per cluster. Programs driven by a single patient, technical artifacts or non-epithelial signal were discarded, and the remaining programs were annotated using histology and IHC.

### Identification of Morphological Motifs by Deep Learning and Non-Negative Matrix Factorization

Whole-slide HES images were tiled at 112 µm, and tile-level gene expression was predicted from UNI v1 features with a multilayer perceptron (MLP). Consensus NMF of this predicted expression matrix yielded morphological motifs, which pathologists annotated after artifactual components were removed. Each tile was assigned its dominant motif. ROI-level motif proportions were the fractions of tiles assigned to each motif, the dominant motif was the most frequent one, and a motif was considered present when it covered more than 15% of tiles.

### Spatial Transcriptomics

Sections, including one normal gastric biopsy, were profiled with the Visium Spatial Gene Expression for FFPE kit (10x Genomics; probe set V1, PN-1000334; Space Ranger v2.0.1), except the two sections in Supplementary Fig. S2B, profiled with the Visium CytAssist FFPE kit (probe set V2, PN-1000520; Space Ranger v3.0.1). Each section was decomposed by Non-negative Spatial Factorization (NSF) (34) with L = 2 to 24, and each RNA program or motif was matched to the NSF factor whose gene loadings correlated best with it. Only IPMN spots were scored, and spots above the median for at least one factor were assigned to the highest-scoring factor motifs), the NSF factor whose gene loadings were most correlated with the program was selected. Only spots histologically annotated as IPMN epithelium were used for scoring, and all other spots are displayed in grey. Spots exceeding the median score of IPMN spots for at least one factor were assigned to the factor with the highest score, and the remaining spots were left uncolored.

### Subgroup Classification Within Low-Grade Intestinal Lesions

Low-grade intestinal lesions were divided into two subgroups by model-based clustering (Mclust) of their *Intestinal Progression* and *Intestinal Shifted* scores.

### Projection onto PDAC Cohorts

IPMN RNA programs and published PDAC subtype signatures (19, 22–26) were projected onto the VST-normalized TCGA-PAAD (18), CPTAC (21), Maurer (20) and Puleo (19) cohorts, and their sample-level scores were compared by Pearson correlation (Fig. 5A).

### Mucus-Based Dysplasia Prediction Model

Each sample was described by ten features: its projection scores on the nine RNA programs, obtained by non-negative least squares (RcppML), and Fanova, a chromosomal imbalance score. Features were z-scored within each cohort. Prediction proceeded in two steps: a phenotype model (intestinal vs non-intestinal), followed by a dysplasia model (high-vs low-grade) specific to the predicted phenotype. Models were trained on tissue RNA-seq (418 samples from 127 patients). After comparing XGBoost and elastic-net logistic regression (glmnet) by 5-fold cross-validation (Supplementary Fig. S5B), we retained XGBoost for the phenotype and intestinal dysplasia models and glmnet for the non-intestinal dysplasia model, with thresholds set by Youden’s index. A unified dysplasia model without phenotype assignment served as a comparator (Supplementary Fig. S5A). The final models were applied without modification to mucus RNA-seq from an independent cohort of 94 patients (56 intestinal, 30 non-intestinal and 8 biphenotypic lesions), and performance was evaluated by ROC AUC, Wilcoxon and Kruskal-Wallis tests (Fig. 5D, 5E).

## Data Availability

Raw sequencing data (FASTQ files) and gene-level count matrices from tissue and mucus RNA sequencing and from spatial transcriptomics are available from the corresponding authors upon request and will be deposited in the Gene Expression Omnibus (GEO) at the request of the journal. The top-ranked genes of each RNA program and morphological motif are provided in Supplementary Tables S1 and S2. Public datasets analyzed in this study were obtained from TCGA-PAAD through the Genomic Data Commons (18), CPTAC through the Proteomic Data Commons (21), ArrayExpress (E-MTAB-6134) (19) and GEO (GSE93326) (20). Code for RNA sequencing processing and RNA-based *KRAS*/*GNAS* variant calling is available at https://github.com/genehetx/RNApipeline (v1.7); all other analysis code is available from the corresponding authors upon request.

## Acknowledgments

This work was supported by the Fondation ARC pour la recherche sur le cancer (IMPULSE: IpMn dysPlasia mULtiscales aSsessment), the Ligue contre le cancer, the GEFLUC, and the integrated cancer research center SiRIC InsiTu “Insights into cancer: From inflammation to tumor” (grant number INCa-DGOS-INSERM-ITMO Cancer_18008).

## Abbreviations

IPMN: intraductal papillary mucinous neoplasm
PDAC: pancreatic ductal adenocarcinoma
MRI: magnetic resonance imaging
CI: confidence interval
PB: pancreatobiliary
IOPN: intraductal oncocytic papillary neoplasm
WHO: World Health Organization
cAMP: cyclic adenosine monophosphate
PPCN: pancreatic papillary cystic neoplasm
PCA: principal component analysis
NMF: non-negative matrix factorization
tVAF: transcribed variant allele frequency
HES: hematoxylin-eosin-saffron
ROI: region of interest
UMAP: uniform manifold approximation and projection
TCGA: The Cancer Genome Atlas
CPTAC: Clinical Proteomic Tumor Analysis Consortium
AUC: area under the ROC curve
PanIN: pancreatic intraepithelial neoplasia
FFPE: formalin-fixed paraffin-embedded
VST: variance-stabilizing transformation
GSEA: gene set enrichment analysis
GESECA: gene set co-regulation analysis
MLP: multilayer perceptron
NSF: non-negative spatial factorization
ROC: receiver operating characteristic
GEO: GEO Gene Expression Omnibus
LG: low-grade
HG: high-grade
IHC: immunohistochemistry

