## Supplemental figures for "Distinct Progression Paths Underlie the Diversity of Pancreatic Papillary Cystic Neoplasms"

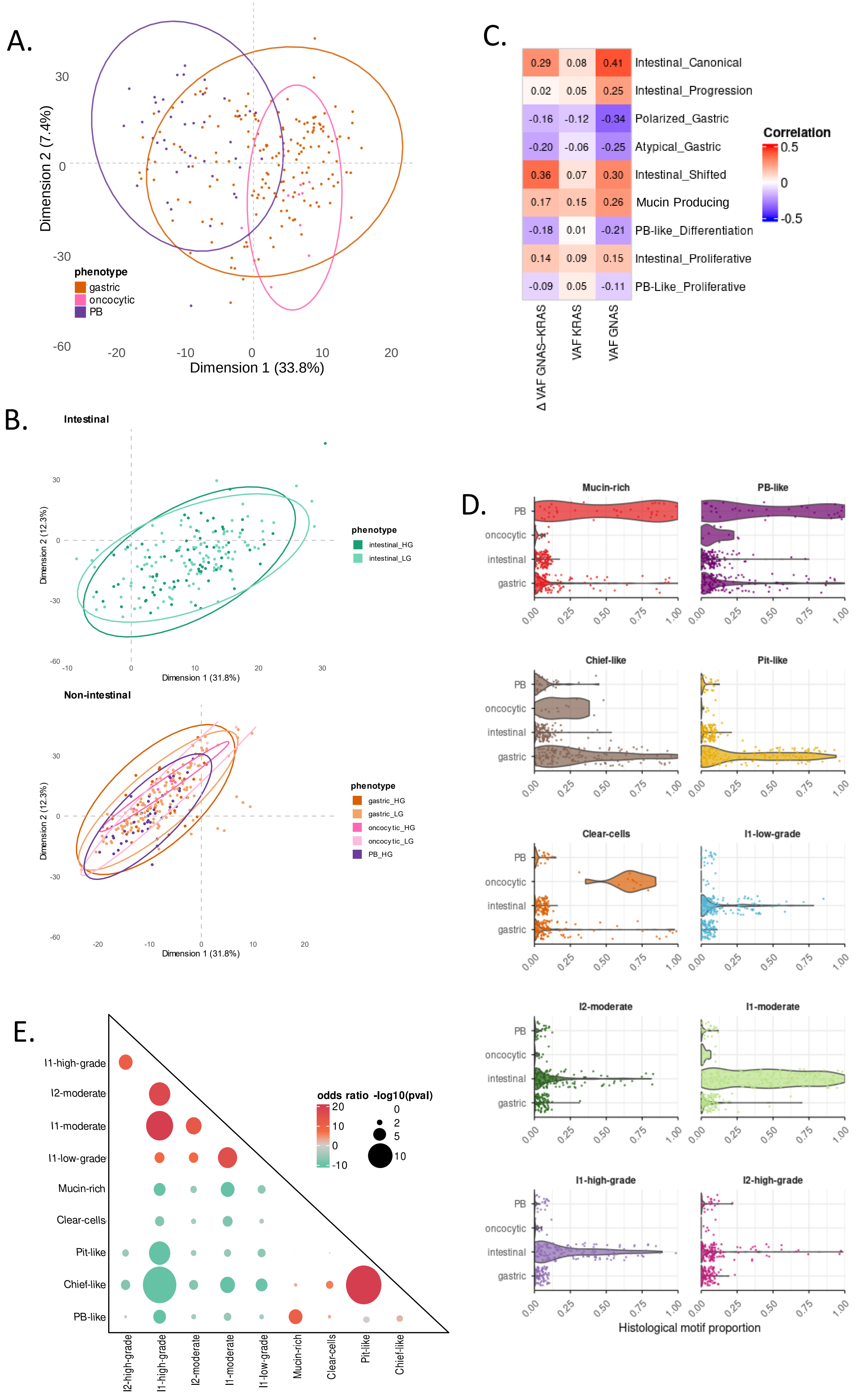
A.
Dimension 2 (7.4%)
Dimension 1 (33.8%)
phenotype
gastric
oncocytic
PB
C.
0.29 0.08 0.41 Intestinal_Canonical
0.02 0.05 0.25 Intestinal_Progression
-0.16 -0.12 -0.34 Polarized_Gastric
-0.20 -0.06 -0.25 Atypical_Gastric
0.36 0.07 0.30 Intestinal_Shifted
0.17 0.15 0.26 Mucin Producing
-0.18 0.01 -0.21 PB-like_Differentiation
0.14 0.09 0.15 Intestinal_Proliferative
-0.09 0.05 -0.11 PB-Like_Proliferative
Δ VAF GNAS−KRAS
VAF KRAS
VAF GNAS
Correlation
B.
Intestinal
Dimension 2 (12.3%)
Dimension 1 (31.8%)
phenotype
intestinal_HG
intestinal_LG
Non-intestinal
gastric_HG
gastric_LG
oncocytic_HG
oncocytic_LG
PB_HG
D.
Mucin-rich
PB-like
Chief-like
Pit-like
Clear-cells
I1-low-grade
I2-moderate
I1-moderate
I1-high-grade
I2-high-grade
PB
oncocytic
intestinal
gastric
Histological motif proportion
E.
I1-high-grade
I2-moderate
I1-moderate
I1-low-grade
Mucin-rich
Clear-cells
Pit-like
Chief-like
PB-like
I2-high-grade
odds ratio
-log10(pval)

**Supplementary Figure 1. Molecular and morphological relationships across IPMN phenotypes (A)** Principal component analysis (PCA) of transcriptomic profiles of gastric, oncocytic and pancreatobiliary IPMN samples. **(B)** PCA of intestinal and non-intestinal IPMN samples, colored by dysplasia grade and phenotype. **(C)** Pearson correlations between KRAS and GNAS mutation status or tVAF (including the difference between *GNAS* and *KRAS* tVAF) and tissue-derived RNA program scores. **(D)** Distribution of morphological motif contributions across oncocytic, gastric/PB and intestinal IPMN phenotypes. **(E)** Co-occurrence of morphological motifs across IPMN samples.

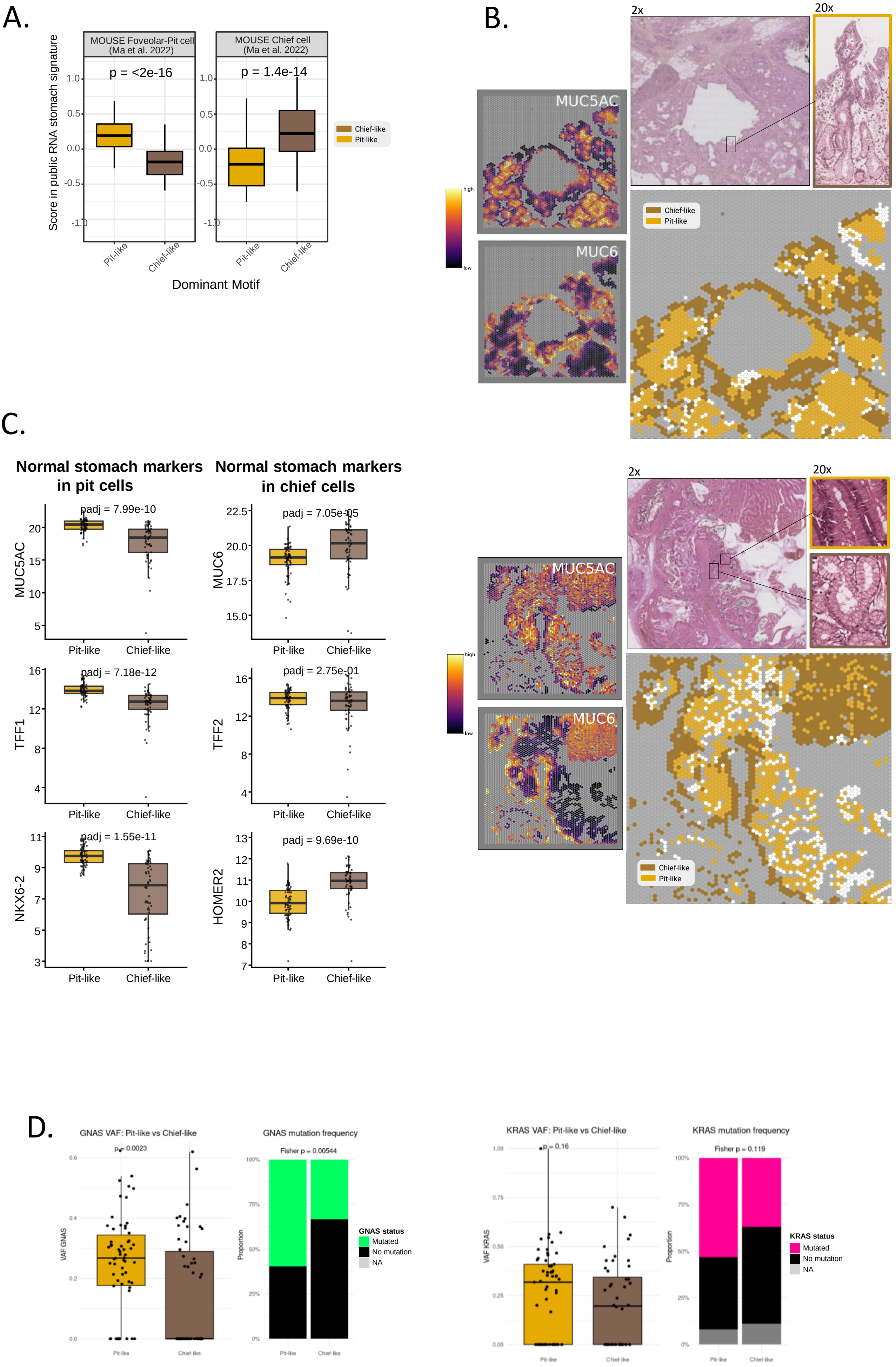
A.
MOUSE Foveolar-Pit cell (Ma et al. 2022)
MOUSE Chief cell (Ma et al. 2022)
p = <2e-16
p = 1.4e-14
Score in public RNA stomach signature
Dominant Motif
Pit-like
Chief-like
B.
2x
20x
MUC5AC
MUC6
high
low
C.
Normal stomach markers in pit cells
Normal stomach markers in chief cells
padj = 7.99e-10
padj = 7.05e-05
padj = 7.18e-12
padj = 2.75e-01
padj = 1.55e-11
padj = 9.69e-10
MUC5AC
MUC6
TFF1
TFF2
NKX6-2
HOMER2
D.
GNAS VAF: Pit-like vs Chief-like
p = 0.0023
VAF GNAS
GNAS mutation frequency
Fisher p = 0.00544
Proportion
GNAS status
Mutated
No mutation
NA
KRAS VAF: Pit-like vs Chief-like
p = 0.16
VAF KRAS
KRAS mutation frequency
Fisher p = 0.119
KRAS status

**Supplementary Figure 2. The gastric axis: epithelial polarity and conservation of gastric epithelial identity. (A)** GESECA enrichment scores for mouse foveolar/pit cell and chief cell reference signatures in Pit-like and Chief-like dominant samples. **(B)** Spatial mapping of Pit-like and Chief-like motifs in two IPMN sections (Visium CytAssist). For each section: HES staining of the capture area (2×), with magnified views (20×) of a pit-cell-enriched and a chief-cell-enriched region; spatial distribution of the NSF factors most correlated with the Pit-like (yellow) and Chief-like (brown) motifs, each IPMN spot being assigned to the factor with the highest score among spots above the median (spots below both thresholds in white); and spatial expression of *MUC5AC* (pit-cell marker) and *MUC6* (chief-cell marker), scaled per gene between the 1st and 99th percentiles of IPMN spots. Non-IPMN spots are shown in grey. **(C)** Expression of canonical normal-stomach pit-cell (*MUC5AC*, *TFF1*, *NKX6-2*) and chief-cell (*MUC6, TFF2, HOMER2*) markers in Pit-like versus Chief-like dominant IPMN samples (Wilcoxon test, BH-adjusted p-values). **(D)** *GNAS* tVAF and mutation frequency in Pit-like versus Chief-like dominant samples, alongside *KRAS* mutation frequency.

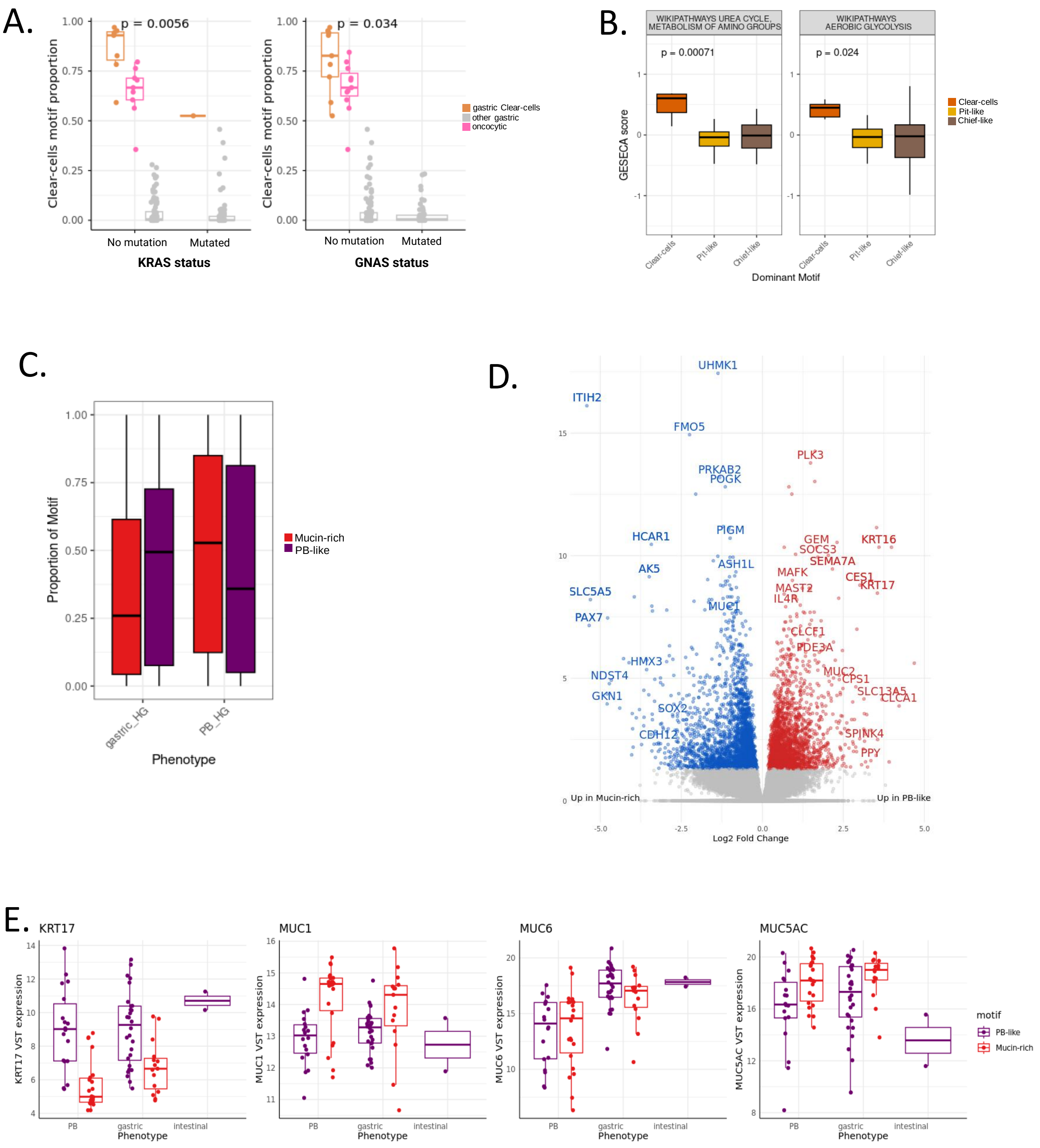

A.
p = 0.0056
Clear-cells motif proportion
1.00
0.75
0.50
0.25
0.00
No mutation
Mutated
KRAS status
p = 0.034
No mutation
Mutated
GNAS status
gastric Clear-cells
other gastric
oncocytic
B.
WIKIPATHWAYS UREA CYCLE, METABOLISM OF AMINO GROUPS
WIKIPATHWAYS AEROBIC GLYCOLYSIS
p = 0.00071
p = 0.024
GESECA score
Clear-cells
Pit-like
Chief-like
Dominant Motif
C.
Proportion of Motif
gastric_HG
PB_HG
Phenotype
Mucin-rich
PB-like
D.
UHMK1
ITIH2
FMO5
PLK3
PRKAB2
POGK
PIGM
HCAR1
GEM
SOCS3
KRT16
SEMA7A
ASH1L
AK5
MAFK
CES1
KRT17
MAST2
SLC5A5
IL4R
MUC1
PAX7
CLCF1
PDE3A
HMX3
NDST4
MUC2
CPS1
SLC13A5
CLCA1
GKN1
SOX2
SPINK4
CDH12
PPY
Up in Mucin-rich
Up in PB-like
Log2 Fold Change
E.
KRT17
KRT17 VST expression
MUC1
MUC1 VST expression
MUC6
MUC6 VST expression
MUC5AC
MUC5AC VST expression
PB
gastric
intestinal
Phenotype
motif
PB-like
Mucin-rich

**Supplementary Figure 3. Diversification of gastric epithelial programs toward oncocytic and PB-associated states. (A)** Proportion of the Clear-cells motif in oncocytic IPMNs, gastric IPMNs enriched for the *Clear-cells* motif, and other gastric IPMNs, stratified by KRAS and GNAS mutation status. **(B)** GESECA enrichment of the urea cycle and glycolysis pathways in *Clear-cells* dominant samples. **(C)** Proportion of PB-like and *Mucin-rich* dominant motifs in PB and high-grade gastric samples. **(D)** Volcano plot of differentially expressed genes between *Pit-like* and *Mucin-rich* dominant samples. **(E)** Expression of *KRT17, MUC1, MUC6* and *MUC5AC* in *PB-like* and *Mucin-rich* dominant samples, stratified by phenotype

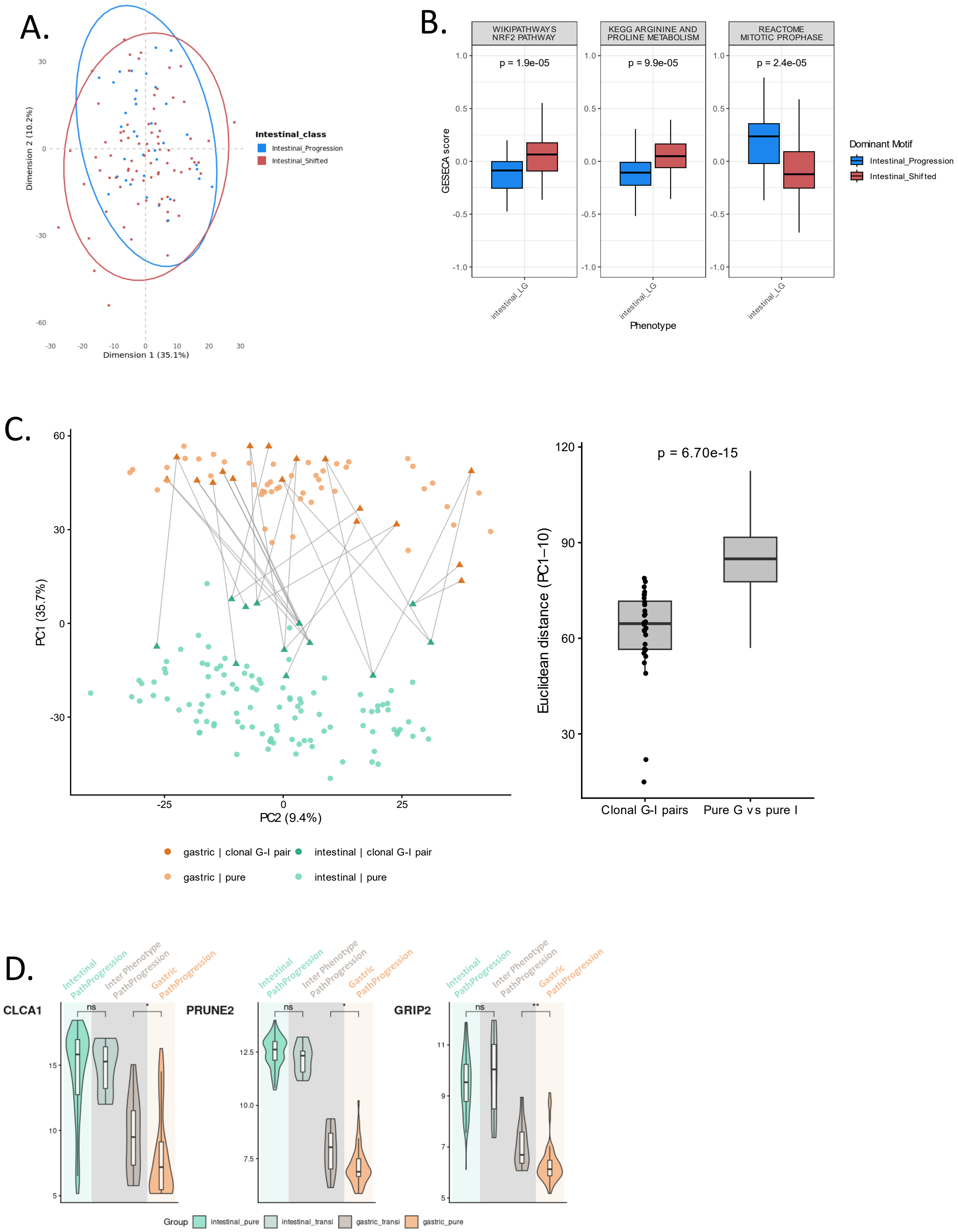
A.
Dimension 2 (10.2%)
Dimension 1 (35.1%)
Intestinal_class
Intestinal_Progression
Intestinal_Shifted
B.
WIKIPATHWAYS NRF2 PATHWAY
KEGG ARGININE AND PROLINE METABOLISM
REACTOME MITOTIC PROPHASE
p = 1.9e-05
p = 9.9e-05
p = 2.4e-05
GESECA score
intestinal_LG
Phenotype
Dominant Motif
Intestinal_Progression
Intestinal_Shifted
C.
PC1 (35.7%)
PC2 (9.4%)
p = 6.70e-15
Euclidean distance (PC1–10)
Clonal G-I pairs
Pure G vs pure I
gastric | clonal G-I pair
intestinal | clonal G-I pair
gastric | pure
intestinal | pure
D.
Intestinal PathProgression
Inter Phenotype PathProgression
Gastric PathProgression
CLCA1
PRUNE2
GRIP2
ns
*
**
Group
intestinal_pure
intestinal_transi
gastric_transi
gastric_pure

E.

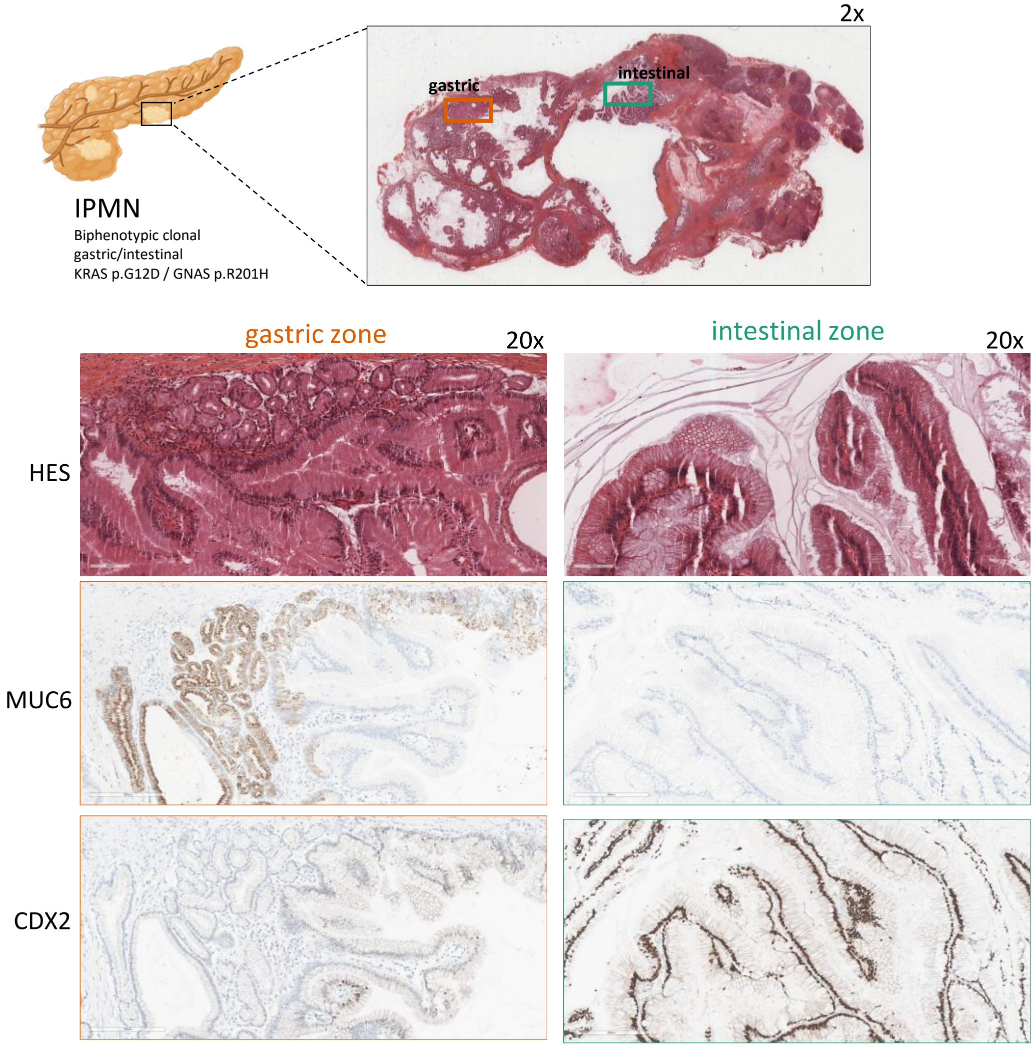
2x
gastric
intestinal
IPMN
Biphenotypic clonal
gastric/intestinal
KRAS p.G12D / GNAS p.R201H
gastric zone
20x
intestinal zone
20x
HES
MUC6
CDX2

F.

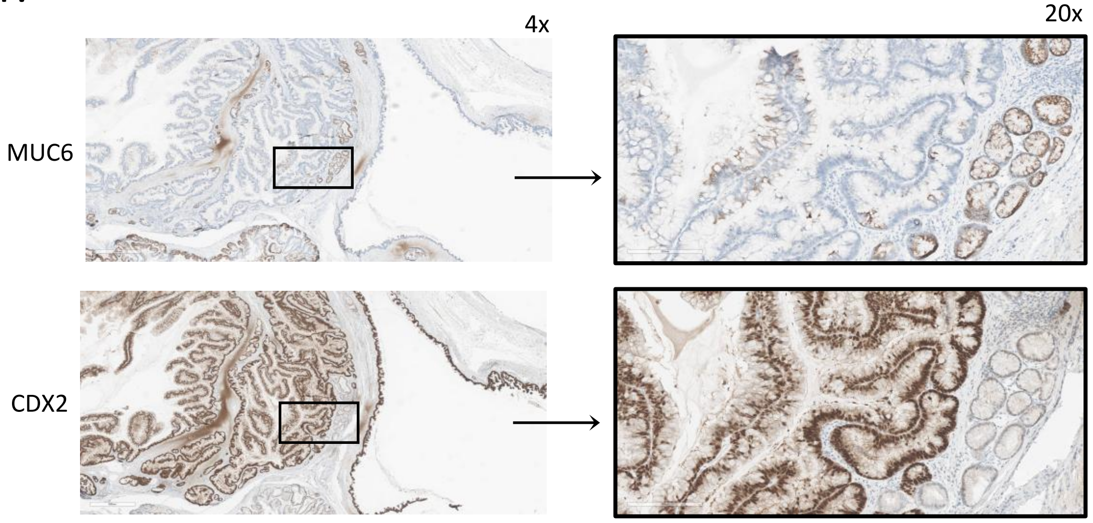
4x
20x
MUC6
CDX2

**Supplementary Figure 4. Molecular features associated with intestinal-shifted IPMN evolution (A)** Principal component analysis (PCA) of intestinal IPMN samples according to *Intestinal Progression* and *Intestinal Shifted* program scores. **(B)** GESECA pathway enrichment analysis comparing *Intestinal Progression* and *Intestinal Shifted* lesions. **(C)** PCA of intestinal and gastric IPMN samples, colored according to clonal relationships. **(D)** Expression of gastric and intestinal lineage-associated markers in clonal and non-clonal gastric/intestinal lesion pairs. **(E)** MUC6 and CDX2 immunostaining in a monoclonal biphenotypic IPMN. **(F)** MUC6 and CDX2 immunostaining at the gastric-intestinal interface.

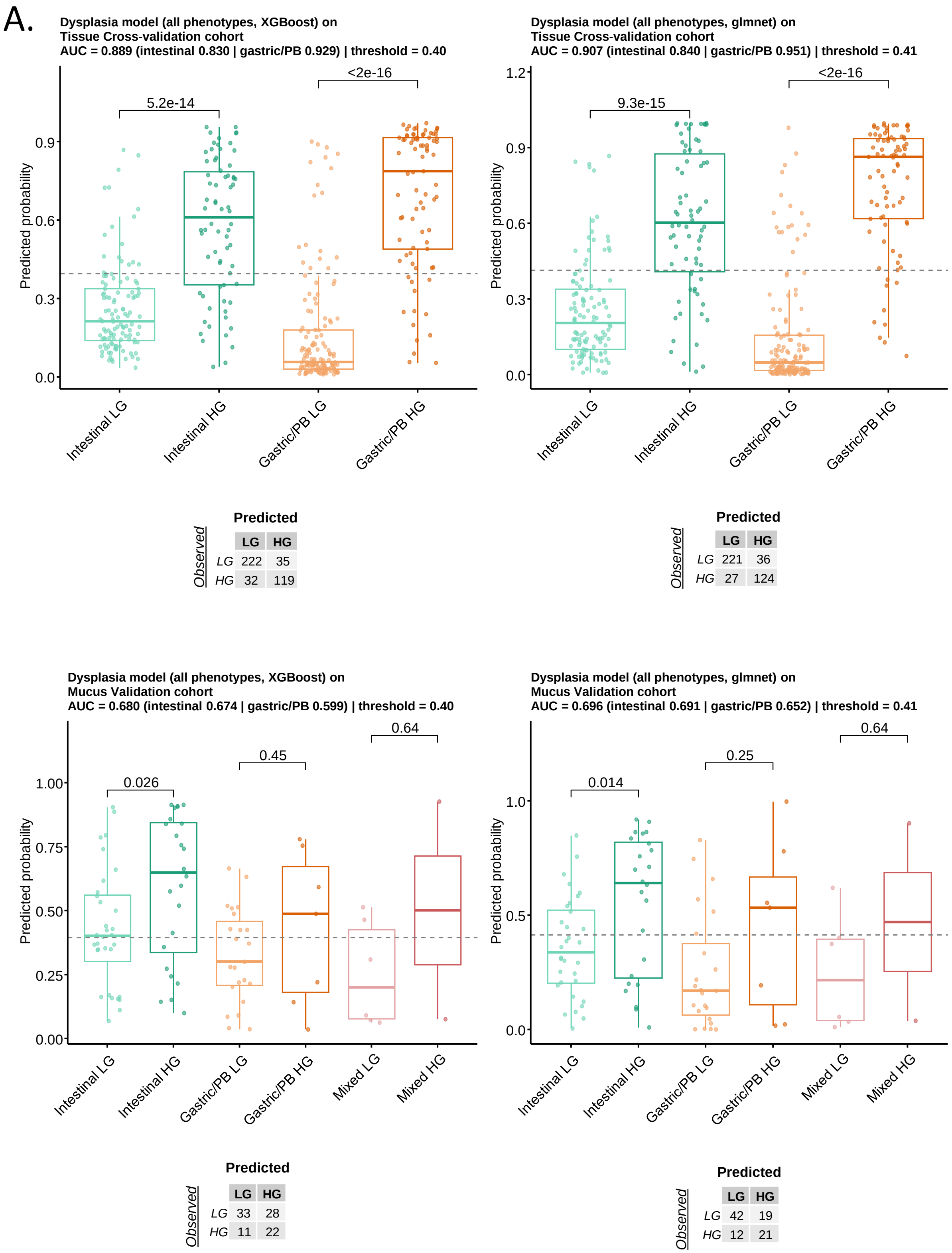
A.
Dysplasia model (all phenotypes, XGBoost) on
Tissue Cross-validation cohort
AUC = 0.889 (intestinal 0.830 | gastric/PB 0.929) | threshold = 0.40
Predicted probability
5.2e-14
<2e-16
Intestinal LG
Intestinal HG
Gastric/PB LG
Gastric/PB HG
Predicted
Observed
LG HG
LG 222 35
HG 32 119
Dysplasia model (all phenotypes, glmnet) on
Tissue Cross-validation cohort
AUC = 0.907 (intestinal 0.840 | gastric/PB 0.951) | threshold = 0.41
9.3e-15
<2e-16
Predicted
Observed
LG HG
LG 221 36
HG 27 124
Dysplasia model (all phenotypes, XGBoost) on
Mucus Validation cohort
AUC = 0.680 (intestinal 0.674 | gastric/PB 0.599) | threshold = 0.40
0.026
0.45
0.64
Mixed LG
Mixed HG
Predicted
Observed
LG HG
LG 33 28
HG 11 22
Dysplasia model (all phenotypes, glmnet) on
Mucus Validation cohort
AUC = 0.696 (intestinal 0.691 | gastric/PB 0.652) | threshold = 0.41
0.014
0.25
0.64
Predicted
Observed
LG HG
LG 42 19
HG 12 21

B.

XGBoost

glmnet

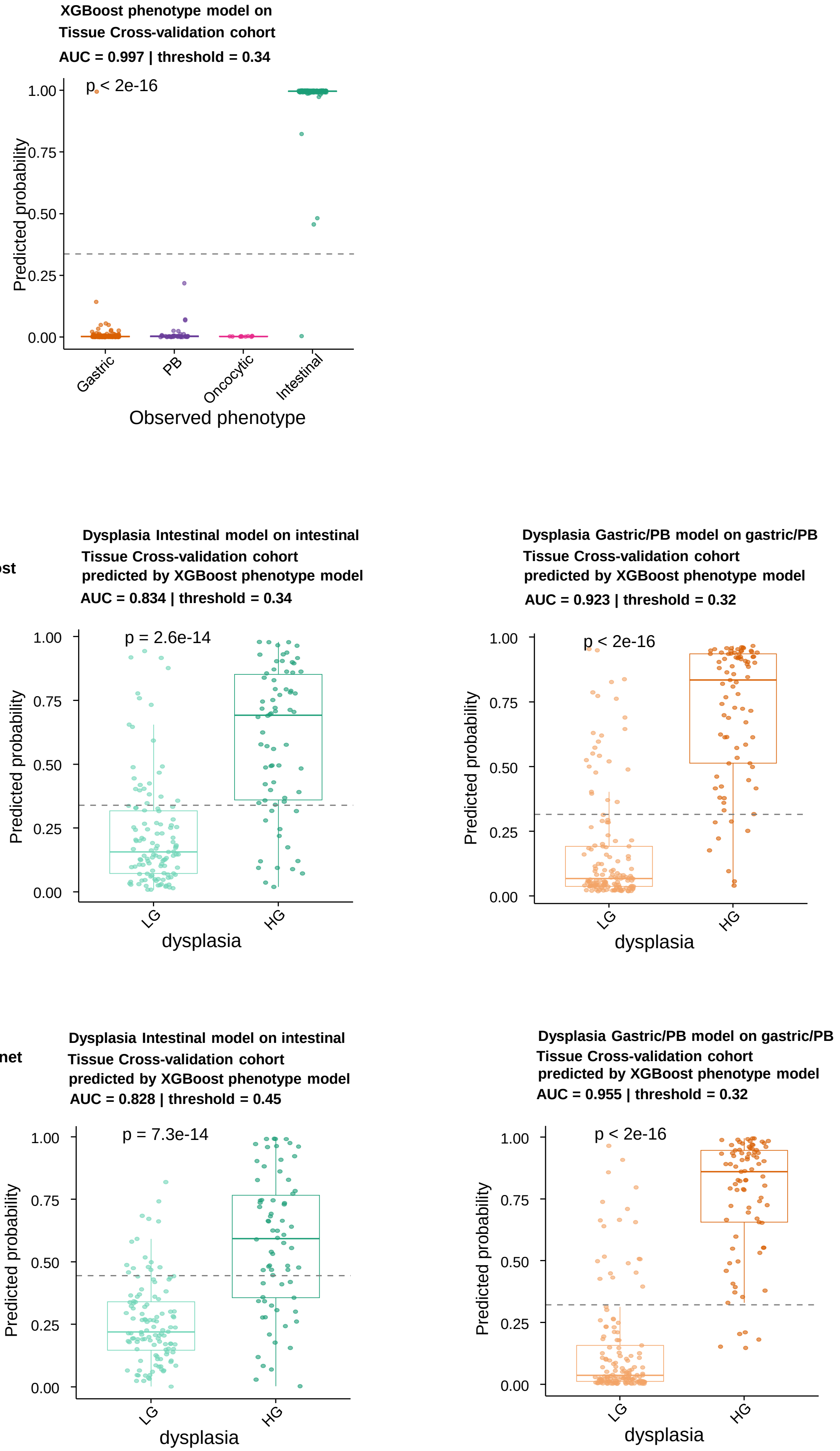

**Supplementary Figure 5. Performance of unified and phenotype-specific dysplasia models. (A)** Unified dysplasia model trained on all ROIs without phenotype assignment. Five-fold cross-validated prediction scores in the tissue cohort and prediction scores in the mucus validation cohort for XGBoost and elastic-net logistic regression (glmnet) models, by observed dysplasia grade and phenotype. Dashed lines indicate Youden-index thresholds. Contingency tables show observed versus predicted classes. p-values: Wilcoxon rank-sum test. **(B)** Five-fold cross-validated prediction scores in the tissue cohort. Top, phenotype model (XGBoost) by observed phenotype. Bottom, intestinal (left) and non-intestinal (gastric/pancreatobiliary, right) dysplasia models trained with XGBoost (upper row) or glmnet (lower row), applied to samples according to the phenotype predicted by the XGBoost phenotype model. Dashed lines indicate Youden-index thresholds. p-values: Kruskal-Wallis (phenotype) and Wilcoxon rank-sum (dysplasia). XGBoost was retained for intestinal dysplasia and glmnet for non-intestinal dysplasia (Fig. 5D, 5E).
