## Supplemental methods for "Distinct Progression Paths Underlie the Diversity of Pancreatic Papillary Cystic Neoplasms"

### Supplementary Methods

##### Patient samples and histological review

The study was approved by the Institutional Review Board (IRB) of the Paris Nord Val-de-Seine University Hospitals Group (HUPNVS) and Paris Cité University (IRB 2024-A02816-41). Surgical pancreatic resections performed for IPMN between 2013 and 2023 were retrieved from the diagnostic database of the Pathology Department at Beaujon Hospital (Clichy, France). All corresponding formalin-fixed, paraffin-embedded (FFPE) and HES slides were reviewed by pancreatic expert pathologists (JC and AC). Histological subtype and dysplasia grade were assessed independently and confirmed through consensus review when necessary. Dysplasia was graded according to the 6th WHO classification: low-grade lesions retained an orderly, mostly single-layered epithelium with basally located, uniform nuclei and preserved polarity, whereas high-grade lesions showed loss of polarity, nuclear pseudostratification, architectural complexity, and marked cytonuclear atypia including nuclear enlargement and irregular nuclear contours. Regions of interest (ROIs) were selected according to IPMN histological subtype (gastric, intestinal, or pancreatobiliary) and grade of dysplasia (Fig. 1), and were chosen so as to exclude any area containing invasive carcinoma. Slides containing the selected ROIs were immunostained with antibodies against mucin 6 (MUC6), mucin 2 (MUC2), mucin 5AC (MUC5AC), and caudal-type homeobox 2 (CDX2) to confirm histopathological subtyping.

##### Histological Staining, Immunohistochemistry, and Digitization

HES staining (3 µm sections) and immunohistochemistry (3 µm sections) were performed on the fully automated ST5020 multistainer (Leica) and BenchMark ULTRA system (Roche Diagnostics), respectively. Stained slides were digitized using the Aperio AT Turbo scanner (Leica). Immunohistochemical markers included MUC6, MUC5AC, MUC2, and CDX2 for subtype assignment, and MUC1 and CPS1 (Hepar1) for phenotypic characterization. These markers were scored semi-quantitatively by a pathologist on a scale of 0 to 3. The Ki67 labeling index was quantified digitally on immunostained slides using QuPath (v0.4). Positive nuclear staining was detected within each ROI using QuPath's pixel classifier, results were aggregated per region.

##### Tissue processing, RNA extraction, and sequencing

FFPE blocks were sectioned at 10 µm and ROI microdissected using the Avenio Millisect System (Roche Diagnostics). RNA was extracted with the Quick-RNA FFPE Kit (Zymo Research), quantified on a NanoDrop One spectrophotometer (Thermo Fisher Scientific), and assessed for quality on a TapeStation 4150 system using the High Sensitivity RNA ScreenTape Assay (Agilent). Libraries were prepared with the SMARTer Stranded Total RNA-Seq Kit Pico Input Mammalian (Takara Bio). For the independent validation series, mucus was aspirated ex vivo from the cystic cavity of freshly resected specimens during macroscopic examination, snap-frozen, and stored at −80 °C; these samples were obtained from 94 patients operated on for a pancreatic precursor cystic neoplasm at the same institution, with no overlap with the tissue cohort, and RNA was extracted with TRIzol. For every mucus sample, phenotype and dysplasia grade were assigned on the corresponding resection specimen and used as ground truth. All RNA samples were sequenced on a NovaSeq X platform. Reads were aligned to the human reference genome (GRCh38, Ensembl release 107) with STAR, and gene-level expression counts were obtained with featureCounts. Counts were normalized with the variance-stabilizing transformation (VST) implemented in DESeq2, and PCA was performed on the 1,000 most variable genes.

##### Fusion detection

*PRKACA* and *PRKACB* rearrangements were assessed as follows. All oncocytic lesions (n = 11) and all gastric lesions with a dominant *Clear-cells* motif (n = 11) were analyzed at the Department of Genetics, Bichat-Claude Bernard Hospital. RNA was extracted from FFPE samples (Maxwell® RSC RNA FFPE,

Promega) and quantified using a Quantus™ Fluorometer (QuantiFluor® RNA System, Promega). Libraries were prepared with the Archer FusionPlex Pan Solid Tumor v2 panel (Integrated DNA Technologies), which targets 137 genes and detects fusions with any partner at the targeted breakpoint regions. They were then sequenced on a NextSeq 500 (Illumina). Reads were aligned to GRCh37/hg19, and fusions were called and annotated with Archer Analysis v7.4.5 (IDT).

**Sample selection and quality control**

Of the resections retrieved and reviewed, IPMN cases from 135 patients met the histological inclusion criteria and were retained for microdissection. A total of 433 ROIs were microdissected and sequenced. Of these, 15 were excluded because fewer than 20,000 genes were detected, a threshold applied uniformly to all libraries. The 418 retained regions, from 127 patients, comprised 191 intestinal (71 high-grade, 120 low-grade), 173 gastric (36 high-grade, 137 low-grade), 43 pancreatobiliary (43 high-grade, 0 low-grade), and 11 oncocytic (5 high-grade, 6 low-grade) regions, for 155 high-grade and 263 low-grade regions overall. No low-grade pancreatobiliary region was recovered, consistent with the current classification in which the pancreatobiliary phenotype is recognized in high-grade lesions only.

**RNA-Based Variant Calling**

Following alignment, read depth at *KRAS* and *GNAS* hotspot positions was estimated using SAMtools depth. tVAFs were computed at these positions using SAMtools mpileup, restricted to known hotspot coordinates. Mutation calls were made using RNApipeline v1.7 (https://github.com/genehetx/RNApipeline) with the following thresholds: a minimum read depth of 10 reads and a minimum variant allele fraction of 0.2 to call a mutation.

**Clonality Inference**

Clonal relationships were inferred for each pair of ROIs from the same patient by comparing *KRAS* and *GNAS* hotspot mutations at the nucleotide level. Pairs with identical *KRAS* and *GNAS* status and at least one shared mutation were classified as clonal. Pairs sharing an identical mutation in one gene, with the other gene mutated in only one of the two ROIs, were also classified as clonal, the additional mutation being considered subclonal. Pairs carrying distinct mutations without any shared mutation were classified as non-clonal. Pairs without any detected mutation, pairs with a mutation detected in only one ROI, and pairs sharing a mutation in one gene but carrying distinct mutations in the other were classified as undetermined. At the patient level, patients with at least one clonal pair were classified as monoclonal (monophenotypic or biphenotypic, depending on the phenotypes of the clonally related ROIs), patients with only non-clonal pairs as multiclonal, and the remaining patients as undetermined.

**Differential Expression and Pathway Analysis**

Differential gene expression between sample groups was performed using DESeq2. Gene set enrichment was assessed with GSEA on gene lists ranked by differential expression, and with GESECA (fgsea R package) on the normalized expression matrix. Sample-level pathway activity scores were computed by averaging mean-centered expression values of pathway genes across the normalized expression matrix. Mouse gastric epithelial reference signatures (16) were converted to their human orthologs before projection.

**Identification of Recurrent RNA Programs by Non-Negative Matrix Factorization**

Transcriptomic profiles were generated from bulk RNA sequencing of microdissected IPMN epithelial regions. Non-negative matrix factorization (NMF) was applied with k ranging from 2 to 30 and random initialization, yielding multiple decompositions. Resulting NMF programs were hierarchically clustered using Jaccard distance, and clusters were defined by maximizing the average silhouette width. A single representative NMF program was selected per cluster based on maximal intra-cluster correlation. Programs were subsequently filtered to exclude programs associated with a single patient, technical biases such as RNA quality artifacts, and non-epithelial signals identified through top gene analysis and pathway enrichment. Retained programs were annotated post hoc using histological labels provided by pathologists and validated through gene expression and immunohistochemical data.

##### Identification of Morphological Motifs by Deep Learning and Non-Negative Matrix Factorization

Digitized slides were tessellated into 112-µm tiles. Tile-level transcriptomic profiles were inferred from HES images using a multilayer perceptron (MLP) trained on UNI v.1 features. For each tile, RNA expression was predicted from histological image features, generating a pseudo-transcriptomic matrix at tile resolution. Consensus NMF was then applied to this matrix to decompose expression profiles into recurrent morphological components. Components associated with image quality artifacts like bubbles and components whose top genes were exclusively non-coding were excluded. Retained components were annotated by pathologists based on their morphological characteristics. Each tile was assigned a dominant morphological motif defined as the component with the highest contribution weight. Motif proportions per ROI were computed as the fraction of tiles assigned to each motif; the dominant motif of an ROI was the motif with the highest proportion, and a motif was considered present in an ROI when its proportion exceeded 15%. Morphological motif score vectors were projected into two-dimensional space using UMAP based on Euclidean distance.

##### Spatial Transcriptomics

FFPE sections were profiled using the Visium Spatial for FFPE Gene Expression kit (10x Genomics; Human Transcriptome, probe V1, PN-1000334; four capture areas per slide, without CytAssist), and reads were mapped with Space Ranger v2.0.1. A normal gastric biopsy, obtained under the same institutional approval, was processed with the same workflow. The two sections shown in Supplementary Figure 2B were processed with the Visium CytAssist Spatial Gene Expression for FFPE kit (Human Transcriptome, probe V2, 6.5 mm, PN-1000520; two capture areas) and mapped with Space Ranger v3.0.1. Spots with fewer than 50 counts and genes detected in fewer than 1% of spots were removed, and counts were normalized by spot size. Each section was analyzed independently using Non-negative Spatial Factorization (NSF; (34)), with models fitted for L = 2 to 24 factors. For each RNA program or motif identified (e.g., *Intestinal Proliferative*, *Intestinal Shifted* and *Polarized Gastric* programs, *Pit-like* and *Chief-like* motifs), the NSF factor whose gene loadings were most correlated with the program was selected. Only spots histologically annotated as IPMN epithelium were used for scoring, and all other spots are displayed in grey. Spots exceeding the median score of IPMN spots for at least one factor were assigned to the factor with the highest score, and the remaining spots were left uncolored.

##### Mucus-Based Dysplasia Prediction Model

Ten features were used: the nine IPMN RNA program projection scores and Fanova, a chromosomal imbalance score computed per sample as the ANOVA F-statistic of gene expression across chromosome arms, averaged. Tissue and mucus RNA-seq data (upper-quartile normalized) were projected onto the tissue-derived NMF programs by non-negative least squares (RcppML), and each feature was z-scored within its cohort. Prediction followed two sequential steps. A phenotype model (intestinal vs. non-intestinal) assigned each sample to a predicted phenotype group according to its threshold. Each group was then scored by a dedicated dysplasia model (high- vs. low-grade). Models were trained on tissue RNA-seq (418 samples from 127 patients) with histopathological phenotype and grade as ground truth. The phenotype model used all samples: 186 intestinal and 222 non-intestinal (170 gastric, 41 pancreatobiliary, 11 oncocytic). Dysplasia models were trained on intestinal (70 high-grade, 116 low-grade) and non-intestinal (81 high-grade, 141 low-grade) lesions, respectively. XGBoost and elastic-net logistic regression (glmnet, $\alpha = 0.5$) were compared by 5-fold cross-validation on tissue, with dysplasia models applied to the predicted phenotype groups (Supplementary Fig. 5B). XGBoost was retained for phenotype and intestinal dysplasia, and glmnet for non-intestinal dysplasia; phenotype models performed equivalently, and XGBoost was kept for consistency. XGBoost hyperparameters were tuned by 5-fold cross-validation, and the glmnet penalty by internal cross-validation. Thresholds were set on cross-validated tissue predictions by Youden's index. For comparison, a unified dysplasia model was trained on all tissue samples without phenotype assignment, using the same features, algorithms and threshold procedure, and applied unchanged to the mucus cohort (Supplementary Fig. S5A).

Final models were trained on the full tissue cohort and applied, with thresholds unchanged, to RNA-seq from mucus aspirated from resected IPMN specimens of an independent cohort of 94 patients. This cohort comprised 56 intestinal (24 high-grade, 32 low-grade), 30 non-intestinal (7 high-grade, 23 low-grade) and 8 mixed-phenotype lesions (2 high-grade, 6 low-grade). Mixed lesions were excluded from phenotype evaluation but scored for dysplasia through their predicted phenotype. Performance was assessed by ROC AUC (pROC), Wilcoxon rank-sum (dysplasia) and Kruskal-Wallis (phenotype) tests, and contingency tables (Fig. 5D-E).

**Statistical analysis**

Statistical comparisons were performed using Fisher's exact test for categorical variables and the Wilcoxon rank-sum test for comparisons between two groups of continuous variables. For comparisons involving more than two groups, the Kruskal-Wallis test was used. Paired comparisons used the Wilcoxon signed-rank test, and correlations were assessed with Pearson's or Spearman's coefficients as indicated. Pairwise comparisons of program scores between phenotypes (Fig. 1D) used Kruskal-Wallis tests. Where indicated, p-values were adjusted for multiple testing with the Benjamini-Hochberg procedure. Unless otherwise specified, tests were two-sided and $p < 0.05$ was considered statistically significant.

**Subgroup Classification Within Low-Grade Intestinal Lesions**

Low-grade intestinal samples were classified into two subgroups based on their *Intestinal Progression* and *Intestinal Shifted* programs scores using model-based clustering implemented in the Mclust R package.

**Projection onto PDAC Cohorts**

Expression data from the TCGA-PAAD (18), CPTAC (21), Maurer (20) and Puleo (19) PDAC cohorts were VST-normalized. IPMN RNA programs and published PDAC subtype signatures (19, 22-26) were projected onto each cohort by non-negative least squares (RcppML). For each dataset, Pearson correlations between the resulting sample-level score (H) matrices yielded a similarity score between each IPMN program and each PDAC signature, displayed as a heatmap (Fig. 5A).

**References**

(Numbering refers to the main-text reference list.)
